# Structure of the microtubule anchoring factor NEDD1 bound to the γ-tubulin ring complex

**DOI:** 10.1101/2024.11.05.622067

**Authors:** Hugo Muñoz-Hernández, Yixin Xu, Daniel Zhang, Allen Xue, Amol Aher, Aitor Pellicer Camardiel, Ellie Walker, Florina Marxer, Tarun M. Kapoor, Michal Wieczorek

## Abstract

The γ-tubulin ring complex (γ-TuRC) is an essential multiprotein assembly, in which γ-tubulin, GCP2-6, actin, MZT1 and MZT2 form an asymmetric cone-shaped structure that provides a template for microtubule nucleation. The γ-TuRC is recruited to microtubule organizing centers (MTOCs), such as centrosomes and pre-existing mitotic spindle microtubules, via the evolutionarily-conserved attachment factor NEDD1. NEDD1 contains an N-terminal WD40 domain that binds to microtubules, and a C-terminal domain that associates with the γ-TuRC. However, the structural basis of the NEDD1-γ-TuRC interaction is not known. Here, we report cryo-electron microscopy (cryo-EM) structures of NEDD1 bound to the human γ-TuRC in the absence or presence of the activating factor CDK5RAP2, which interacts with GCP2 to induce conformational changes in the γ-TuRC and promote its microtubule nucleating function. We found that the C-terminus of NEDD1 forms a tetrameric α-helical assembly that contacts the lumen of the γ-TuRC cone, is anchored to GCP4, 5 and 6 via protein modules consisting of MZT1 & GCP3 subcomplexes, and orients its microtubule-binding WD40 domains away from the complex. We biochemically tested our structural models by identifying NEDD1 mutants unable to pull-down *γ*-tubulin from cultured cells. The structure of the γ-TuRC simultaneously bound to NEDD1 and CDK5RAP2 reveals that both factors can associate with the “open” conformation of the complex. Our results show that NEDD1 does not induce conformational changes in the γ-TuRC, but suggest that anchoring of γ-TuRC-capped microtubules by NEDD1 would be structurally compatible with the significant conformational changes experienced by the γ-TuRC during microtubule nucleation.

## Introduction

Microtubules, cytoskeletal filaments found in all eukaryotic cells, are nucleated by the γ-tubulin ring complex (γ-TuRC). This ∼2.3 MDa complex forms a template that helps arrange α/β-tubulin into the canonical 13-protofilament, pseudo-helical microtubule lattice architecture (Aher et al., 2024; Brito et al., 2024; Dendooven et al., 2024). The γ-TuRC is composed of the core structural subunits γ-tubulin, γ-tubulin complex protein 2 (GCP2), GCP3, GCP4, GCP5, and GCP6, along with mitotic spindle organizing proteins associated with a ring of γ-tubulin 1 (MZT1) and MZT2 (Kollman et al., 2011). These proteins are arranged into an asymmetric cone-shaped structure containing 14 “spokes”, each of which comprises a γ-tubulin molecule supported by a GCP2/3/4/5/6 subunit. The asymmetry of the γ-TuRC stems from the unique arrangement of GCP subunits in the complex, where: positions 1-8 of the γ-TuRC contain an alternating array of GCP2 and GCP3; GCP4, GCP5, GCP4, and GCP6 then occupy positions 9-12; and the last two positions are occupied again by GCP2 and GCP3 (Wieczorek et al., 2020b; Liu et al., 2019; Consolati et al., 2020; Zimmermann et al., 2020). The terminal GCP3 subunit at position 14 overlaps with GCP2 at position 1, accommodating the heterotypic α-tubulin:β-tubulin lateral contacts formed at the microtubule “seam” (Aher et al., 2024; Brito et al., 2024; Dendooven et al., 2024). It has been proposed that the arrangement of GCP4-GCP6 would present unique protein-binding interfaces for γ-TuRC regulators (Wieczorek et al., 2020b), but binding partners recognizing this particular subunit configuration in the γ-TuRC have not yet been characterized.

Two well-documented factors that associate with the γ-TuRC to regulate its cellular functions are CDK5RAP2 (Choi et al., 2010), whose conserved CM1 motif binds to the γ-TuRC to induce conformational changes that promote microtubule nucleation (Xu et al., 2024; Serna et al., 2024), and NEDD1 (neural precursor cell expressed, developmentally down-regulated 1). NEDD1 is a γ-TuRC-associated, cell cycle-regulated protein that is important for γ-TuRC localization and microtubule anchoring at centrosomes and within mitotic spindles (Lüders et al., 2006; Haren et al., 2006). Upregulation of NEDD1 has also been linked to the progression of numerous types of solid tumors, making it a potential therapeutic target (Zhuo et al., 2024). Like CDK5RAP2, NEDD1 is not needed for γ-TuRC assembly. Instead, current models suggest that NEDD1 serves two primary functions. One is to recruit the γ-TuRC to centrosomes in undifferentiated cells (Muroyama et al., 2016), consistent with a role in the nucleation and anchoring of microtubules. The other is to promote microtubule-mediated microtubule nucleation in concert with the Augmin complex in a process also termed “branching” nucleation (Liu and Wiese, 2008; Uehara et al., 2009), which contributes to microtubule assembly in dividing cells (Uehara et al., 2009; Goshima et al., 2008), neurons Cunha-Ferreira et al., 2018; Viais et al., 2021), and plants (Hotta et al., 2012; Liu et al., 2014). Secondary structure predictions suggest that NEDD1 consists of two domains: (1) an N-terminal WD40 repeat domain comprising residues ∼1-301, and (2) a C-terminal α-helical region (residues ∼555-660) (Figure 1A) (Manning et al., 2010). The WD40 domain is characterized by seven tandem repeats that fold into a β-propeller structure, which can bind to microtubules with weak affinity (Zhang et al., 2022). In contrast, NEDD1’s C-terminal region is predicted to be comprised of α-helices (Figure 1A), forms tetramers *in vitro* (Manning et al., 2010), and is responsible for centrosomal localization of the γ-TuRC (Lüders et al., 2006; Haren et al., 2006). Despite the importance of NEDD1 in overall γ-TuRC function, the molecular basis for how it interacts with the γ-TuRC is not known. Further, while recent studies have shown that CDK5RAP2 transitions the γ-TuRC from an open, inactive state into a partially-closed conformation with substantially improved microtubule nucleating activity (Xu et al., 2024; Serna et al., 2024), it is not clear whether NEDD1 alters the conformation of the γ-TuRC, nor whether NEDD1 binding would be compatible with the substantial conformational changes experienced by the γ-TuRC during microtubule nucleation (Aher et al., 2024; Brito et al., 2024).

**Figure 1.**
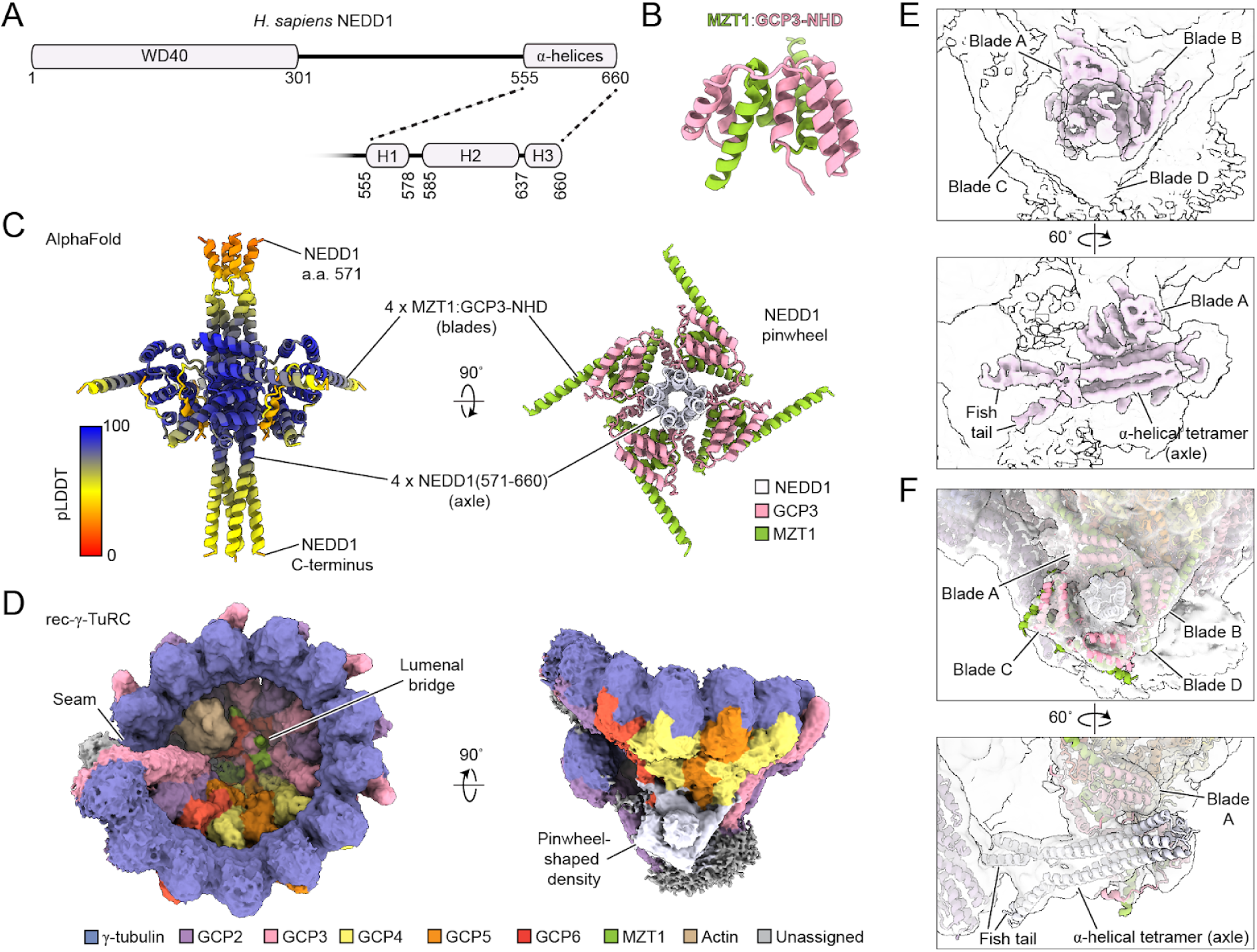
A pinwheel-shaped structure consisting of a tetrameric NEDD1 helical bundle and four MZT1:GCP3-NHD modules docks onto the base of the asymmetric cone-shaped human γ-TuRC. A) Schematic of the human NEDD1 sequence. A zoom-in on the C-terminal helical region predicted to form several α-helices is shown. Secondary structure predictions are taken from UniProt (UniProt Consortium, 2023). B) Cartoon representation of the MZT1:GCP3-NHD structure in the γ-TuRC lumenal bridge, from PDB ID: 6X0U (Wieczorek et al., 2020a). C) Two views of an AlphaFold prediction containing 4 copies each of NEDD1 residues 571-660, MZT1, and GCP3 residues 1-120. The model on the left is coloured according to the predicted local distance difference test (pLDDT) score from the AlphaFold prediction. D) Two views of the consensus recombinant γ-TuRC density map (surface representation). The seam, lumenal bridge, and pinwheel-shaped densities are labeled; the pinwheel-shaped density is coloured in lavender. Map resolution is 4.7 Å, but is shown at a low threshold to include features with weaker density. Higher resolution features can be found in Supplementary Figure S3B. E) Two views of the pinwheel density post-processed using EMready (He et al., 2023) (light pink surface representation). The CryoSPARC post-processed map is shown at the same threshold as in D) as a transparent white surface for reference. The pinwheel axle containing the fish tail and α-helical tetramer, as well as pinwheel blades A-D, are indicated. F) Two views of the refined γ-TuRC model in the rec-γ-TuRC consensus map, focusing on the pinwheel density. Pinwheel features are labeled as in E).

Previous cell biology studies have suggested that the C-terminus of NEDD1 interacts with a portion of GCP3 that includes its “N-terminal α-helical domain” (GCP3-NHD) (Zhang et al., 2022). GCP3-NHD is structurally homologous to the NHDs of GCP2, GCP5, and GCP6, all of which have been found to form subcomplexes together with either MZT1 (GCP3-, GCP5-, or GCP6-NHD) or MZT2 (GCP2-NHD). Different MZT:GCP-NHD subcomplexes constitute protein “modules” that dock across the γ-TuRC to mediate complex assembly and activation (Wieczorek et al., 2020a; Huang et al., 2020; Xu et al., 2024; Serna et al., 2024; Würtz et al., 2022) (Figure 1B). Intriguingly, based on immunoprecipitation and immunofluorescence imaging, MZT1 has also been proposed to mediate the interaction between the γ-TuRC and NEDD1 via GCP3’s N-terminus (Cota et al., 2017; Zhang et al., 2022). There is therefore compelling evidence that NEDD1 interacts with the γ-TuRC via MZT1:GCP3-NHD modules, but the structural basis of these potential interactions are not known.

Here, we report cryo-electron microscopy (cryo-EM) structures of NEDD1 bound to the human γ-TuRC. We found that NEDD1’s C-terminus forms a tetrameric α-helical bundle that interfaces with several core γ-TuRC components, including GCP4-6. The NEDD1 tetramer forms the basis of a pinwheel-shaped structure that is encircled by multiple MZT1:GCP3-NHD subcomplexes. Mutations in NEDD1 predicted to destabilize the pinwheel structure impaired NEDD1-dependent affinity-purification of γ-tubulin complexes from cells. The structure of NEDD1 allowed us to assign previously missing features in the γ-TuRC, including a MZT1-and GCP5-containing subcomplex at the γ-TuRC seam that may stabilize α/β-tubulin dimers during early stages of microtubule nucleation (Brito et al., 2024). Moreover, by determining the structure of the γ-TuRC bound to both NEDD1 and an activating fragment of CDK5RAP2, we demonstrate that both regulatory factors can simultaneously associate with the “open” conformation of the complex. Our results explain how NEDD1 binds the γ-TuRC, and offer insights into the role of distinct regulatory protein binding sites in γ-TuRC recruitment at different microtubule organizing centers (MTOCs).

## Results

To elucidate the structure of NEDD1, as well as how it interacts with the γ-TuRC, we first used AlphaFold to predict potential subcomplexes between NEDD1 and MZT1:GCP3-NHD (Abramson et al., 2024). We found that four copies of the C-terminal residues 571-660 of NEDD1 together with four copies each of MZT1 and residues 1-120 of GCP3 resulted in a high-confidence, “pinwheel”-shaped model, in which four MZT1:GCP3-NHD modules encircle a NEDD1 tetramer at ∼90 degrees relative to one another (hereafter, the “NEDD1 pinwheel”) (Figure 1C and Supplementary Figure S2A-B). The “axle” of the NEDD1 pinwheel comprises four copies of NEDD1 α-helix H2 organized into a four-helix bundle (Figure 1C), which is consistent with the finding that NEDD1 C-terminal residues 572-660 form a tetrameric α-helical assembly *in vitro* (Manning et al., 2010). Four MZT1:GCP3-NHD modules are predicted to surround the axle to form the “blades” of the NEDD1 pinwheel structure (Figure 1C), consistent with the reported interactions between MZT1, GCP3-NHD, and NEDD1 (Cota et al., 2017; Zhang et al., 2022). For comparison, we also performed an AlphaFold prediction for four copies of MZT2:GCP2-NHD with NEDD1 (Supplementary Figure S1C-D), but this model had significantly lower prediction confidence scores compared to the one using MZT1:GCP3-NHD (Supplementary Figure S1A-B). These observations suggest that NEDD1 and MZT1:GCP3-NHD form a hetero-dodecameric subcomplex that may be associated with the γ-TuRC.

To test this hypothesis, we determined the cryo-EM structure of a NEDD1-containing, reconstituted human γ-TuRC ("rec-γ-TuRC") (Wieczorek et al., 2021). In previous studies, we generated several low-to medium-resolution 3D reconstructions of rec-γ-TuRC (Wieczorek et al., 2021; Aher et al., 2024), and consistently observed unassigned densities at the bottom of the cone-shaped assembly (Figure 1D). Notably, such densities have not been reported in other γ-TuRC structures, which used γ-TuRC specimens prepared with little or no NEDD1 (Wieczorek et al., 2020b; Liu et al., 2019; Zimmermann et al., 2020; Consolati et al., 2020; Xu et al., 2024; Würtz et al., 2022; Dendooven et al., 2024; Vermeulen et al., 2024), making them reasonable candidates for a NEDD1 assignment.

To improve the resolution of rec-γ-TuRC as well as to select for resolvable features in the previously unassigned densities, we analyzed a dataset of “free” rec-γ-TuRC present in micrographs collected during a recent cryo-EM study of rec-γ-TuRC-capped microtubule end structures (Aher et al., 2024). We developed a new processing pipeline, in which rec-γ-TuRC particles were re-picked from raw micrographs and subjected to particle clean up in RELION (Kimanius et al., 2021), followed by supervised 3D classifications and refinements in CryoSPARC (Punjani et al., 2017). This targeted procedure improved the resolution of the rec-γ-TuRC reconstruction from ∼7 Å to 4.7 Å (Figure 1D, Supplementary Figure S2A-B, S2E-F, S3A-B and Supplementary Table 2). It also helped resolve the structure of the portion of the unassigned densities associated with GCP4-6 at γ-TuRC positions 9-12, which, at low thresholds, resemble the four-bladed NEDD1 pinwheel model predicted by AlphaFold (Figure 1D). Consequently, the NEDD1 pinwheel model could be rigid body-fitted into this density (Supplementary Figure S3C), suggesting that NEDD1 binds to the γ-TuRC at the bottom of the cone-shaped complex.

The fitted NEDD1-containing AlphaFold model agreed well with the pinwheel density, especially in the axle and Blades A & B (Supplementary Figure S3C). Although the tertiary structure of the two lower MZT1:GCP3-NHD subunits (Blades “C” & “D”) also agreed well with the model at low density thresholds, their relatively weaker density compared with Blades A & B made it difficult to assess their secondary structure elements. MZT1:GCP5-NHD, MZT1:GCP6-NHD, and MZT2:GCP2-NHD all have qualitatively similar folds as MZT1:GCP3-NHD (Wieczorek et al., 2020a; Huang et al., 2020), meaning they could theoretically constitute one or more of the blade densities. However, the following lines of evidence support the exclusive assignment of MZT1:GCP3-NHD to the NEDD1 pinwheel. First, MZT1:GCP6-NHD can be ruled out, since the only copy of GCP6 is found in the lumenal bridge as an actin-binding domain (Figure 1D) (Wieczorek et al., 2020a; Liu et al., 2019; Consolati et al., 2020; Zimmermann et al., 2020; Xu et al., 2024; Würtz et al., 2022; Wieczorek et al., 2021). Second, MZT2 is missing in organisms that retain NEDD1, such as in *D. melanogaster* (Tovey and Conduit, 2018), and, as mentioned, MZT2:GCP2-NHD is not predicted to form a complex with NEDD1 (Supplementary Figure S1C-D). Third, NEDD1 is conserved in flowering plants (Zeng et al., 2009), and *A. thaliana* NEDD1 is predicted to interact with MZT1:GCP3-NHD (Supplementary Figure S1E-F), but not with, e.g., MZT1 and N-terminal portion of GCP5 (Supplementary Figure S1G-H). Lastly, a recent deep learning framework for identifying protein-protein interactors across the human genome scored GCP3 significantly higher than any other GCP subunit as a likely NEDD1 interactor (Zhang et al., 2024). Altogether, these observations argue that the pinwheel density contains four copies each of NEDD1, MZT1, and GCP3-NHD.

### The NEDD1 pinwheel associates with the γ-TuRC through conserved interfaces

Next, we used the density maps and predicted models above to build a molecular model of the human γ-TuRC bound to the NEDD1 pinwheel (Figure 1F, Supplementary Figure S3B and Supplementary Tables S2-S3). Analysis of the NEDD1 pinwheel model highlighted that the interfaces between NEDD1 and the MZT1:GCP3-NHD modules are conserved (Supplementary Figure S1I-J). We identified several potential interfaces that appear to contribute to pinwheel formation and γ-TuRC-docking. Non-polar residues within the NEDD1 tetramer, such as F603 and F622, form hydrophobic cores that may be critical for the formation and stability of the NEDD1 helical assembly (Figure 2A-B). Likewise, electrostatic interactions between NEDD1 residues E598, D602, and E605 and GCP3-NHD residues K57 and K60 likely stabilize the NEDD1-GCP3-NHD interface (Figure 2C-D). Mutating either of these residue sets to alanine (F603A/F622A and E598A/D602A/E605A), which would be predicted to destabilize the NEDD1 tetramer and the NEDD1-MZT1:GCP3-NHD interface, respectively, both reduced the ability of overexpressed NEDD1 to co-immunoprecipitate γ-tubulin from HEK293T cells (Figure 2E). These results support our γ-TuRC-NEDD1 pinwheel model and reveal at least two key interfaces important for facilitating NEDD1-γ-TuRC interactions.

**Figure 2.**
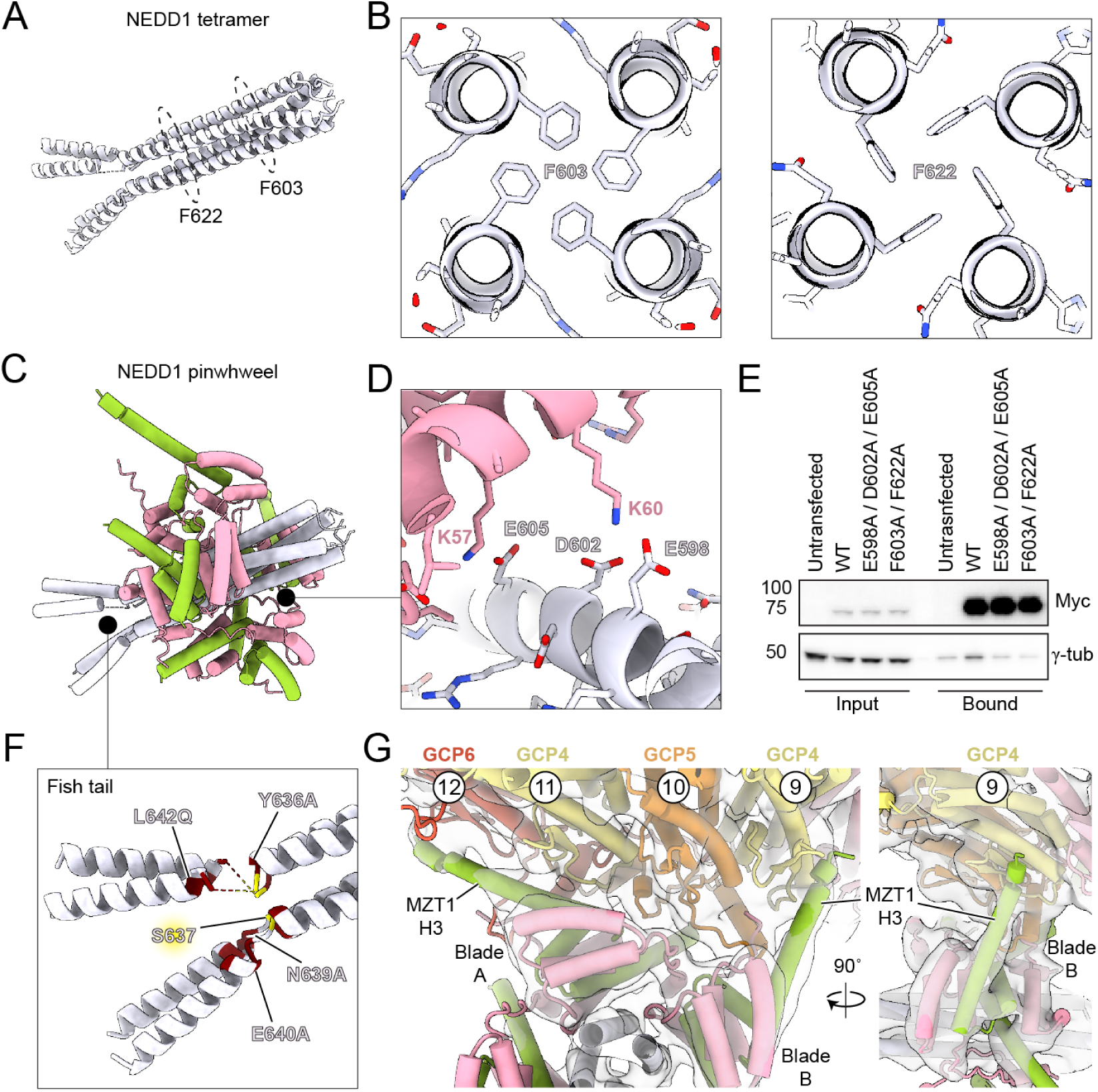
The NEDD1 pinwheel associates with the γ-TuRC through conserved interfaces. A) Cartoon representation of a model of the NEDD1 C-terminal tetramer. Locations of conserved F603 and F622 residues are highlighted by dashed circles. B) Cross-section views of the NEDD1 tetramer AlphaFold model showing the predicted packing of F603 (left) and F622 (right). C) Cartoon representation of a model of the NEDD1 pinwheel. Black circles indicate zoom-in areas of interest for panels D) and E), color legend as in Figure 1F. D) Zoom-in view of NEDD1 pinwheel AlphaFold model for regions specified in C), showing conserved residues involved in electrostatic interactions between NEDD1 and GCP3 in the pinwheel. E) Western blot of inputs and bound fractions of SBP pulldowns of Myc-SBP-NEDD1 constructs from HEK293T. Untransfected cells served as a negative control. The experiment was performed three times with similar results. F) Zoom-in view of the fish-tail region of the NEDD1 pinwheel model, highlighting previously-reported mutations that abolish NEDD1:γ-TuRC interactions (maroon) (Manning et al., 2010), as well as an identified Plk1 phosphorylation site (yellow) (Zhang et al., 2009). G) Two views of the NEDD1 pinwheel Blades A and B bound to the base of the γ-TuRC (cartoon representation with cryo-EM density in transparent grey surface).

Next, we examined the interaction between the NEDD1 pinwheel and the γ-TuRC, which is also mediated by two main interfaces. In the first interface, NEDD1 pinwheel Blades A & B directly contact the underside of the γ-TuRC (Figure 2G). Blade B mainly contacts the GRIP1 domain of GCP4 at position 9, while Blade A forms an extended interface with the GRIP1 domains of GCP5, GCP4, and GCP6 (Figure 2G). In Blade B, MZT1’s C-terminal α-helix (H3) reaches up along GCP4 to terminate nearly halfway up the outer face of its GRIP1 domain (Figure 2G). Additionally, a short α-helical element formed by GCP5 residues ∼243-263 is inserted into a pocket lined by MZT1 α-helices H2 and H3 (Supplementary Figure S3E). In Blade A, MZT1 α-helix H3 traces across the bottom of the GRIP1 domains of GCP4 and GCP6 (Figure 2G). Analogous to Blade B, a short α-helix formed by GCP6 residues ∼325-343 also inserts into a pocket lined by MZT1 H2 and H3 (Supplementary Figure S3E). These observations reveal how the pinwheel blades interface with the γ-TuRC, and suggest that the GCP5 and GCP6-specific GRIP1 insertion elements -which have previously been modeled, but whose purpose was not described (Würtz et al., 2022; Zimmermann et al., 2020) -probably function to dock and orient the NEDD1 pinwheel.

In the other NEDD1 pinwheel-γ-TuRC interface, the C-terminal portion of the NEDD1 tetrameric α-helical bundle density (made of NEDD1 α-helices H2) splays apart into two helical pairs (made of NEDD1 α-helices H3). Notably, the predicted loop separating NEDD1 α-helices H2 and H3 is also the location of previously-reported mutations in NEDD1 that disrupt its ability to co-immunoprecipitate with γ-tubulin (Manning et al., 2010) (Figure 2F). The splayed H3 α-helices, which we term the NEDD1 “fish tail” (Figure 1E-F and 2F), contact the GRIP1 domains of GCP2 at position 1 and GCP3 at position 2 via the top helical pair; the bottom pair appears unattached and, likely as a result, is less well-resolved (Figure 3A and Supplementary Figure S3B). Moreover, the fish-tail:GCP2/3 connection is not direct, but is facilitated by an intermediate loop density that extends from the GCP6 “belt” (Wieczorek et al., 2020a) (Figure 3A; top). An AlphaFold 3 prediction of residues 191-252 adjacent to the GCP6 belt together with the GRIP1 domains of GCP2 and GCP3 resulted in a model that agrees well with our density (Figure 3A and Supplementary Figure S3F). The prediction allowed us to model an extension of the GCP6 belt along the lumenal face of GCP2 and GCP3, which ultimately forms the interface with NEDD1’s C-terminus via the upper helical pair of the fish tail feature (Figure 3A). Together, these observations improve previous models for GCP6 and show that NEDD1 binds to the γ-TuRC directly via its C-terminal residues, as well as indirectly via two MZT1:GCP3-NHD domains.

**Figure 3.**
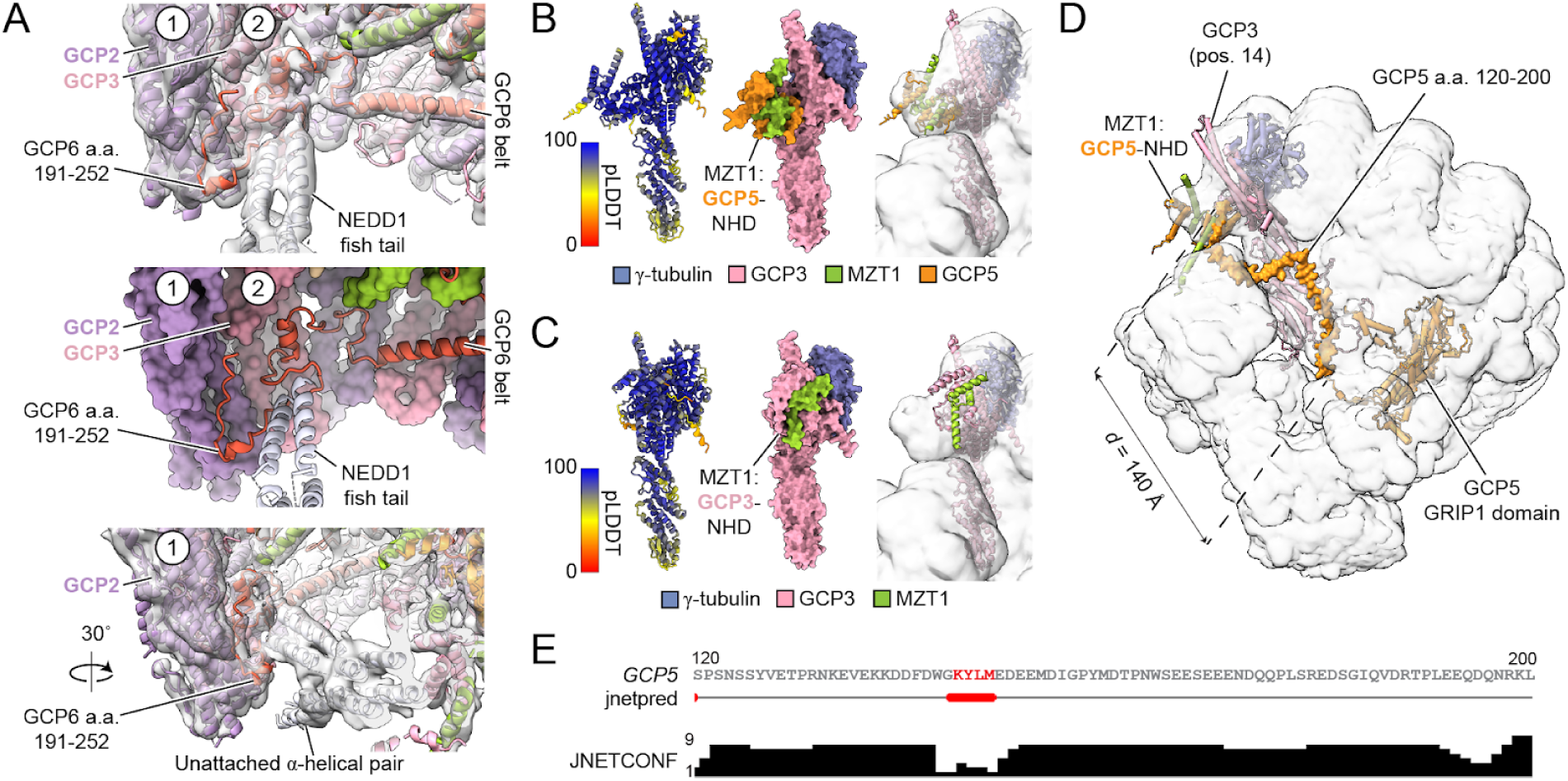
The NEDD1 pinwheel contacts an extension in the GCP6 belt and enables the assignment of MZT1:GCP5-NHD to the γ-TuRC seam. A) Two views of the upper helical pair of the NEDD1 fish tail contacting positions 1 and 2 of the γ-TuRC (top: cartoon representation and cryo-EM density in transparent grey surface; middle: cartoon representation of NEDD1 and GCP6 next to a surface representation of the γ-TuRC). Newly-modeled GCP6 residues 191-252 extending from the GCP6 belt are indicated. The bottom panel shows a rotated view of the same interface to highlight the unattached helical pair in the fish tail. γ-TuRC subunit positions 1 (GCP2) and 2 (GCP3) are indicated, where possible. B) AlphaFold model of GCP3, MZT1:GCP5-NHD, and γ-tubulin. The model on the left is coloured according to the predicted local distance difference test (pLDDT) score from the AlphaFold prediction (cartoon representation). The model on the right is shown in surface representation and is coloured according to the legend. C) AlphaFold model of GCP3, MZT1:GCP3-NHD, and γ-tubulin. The model on the left is coloured according to the predicted local distance difference test (pLDDT) score from the AlphaFold prediction (cartoon representation). The model on the right is shown in surface representation and is coloured according to the legend. D) Surface representation of rigid body-fitted AlphaFold model of GCP5, GCP4, GCP6, GCP2, GCP3, and MZT1 in the rec-γ-TuRC consensus map (transparent representation). MZT1:GCP5-NHD, GCP3, and the disordered GCP5 linker (a.a. 120-200) are indicated. The Euclidean distance between GCP5 residue 120 and 200 in the model is indicated. E) Secondary structure prediction of human GCP5 residues 120-200. Top: primary sequence (red = predicted α-helices); middle: jnetpred secondary structure prediction result (red = α-helices); bottom: confidence score for the prediction. Figure generated using Jalview (Waterhouse et al., 2009).

### The structure of the NEDD1-bound γ-TuRC enables assignment of GCP5-specific features

In our new reconstruction, we also noticed an unassigned α-helical element lining the lumenal face of GCP6’s GRIP1 domain (Supplementary Figure S3G). This feature has been observed in previous γ-TuRC reconstructions as part of a set of densities spanning from γ-TuRC positions 11-14 (Zimmermann et al., 2020; Xu et al., 2024). Using AlphaFold, we found that residues ∼567-608 of GCP5 corresponding to a GCP5-specific insertion element are predicted to form a helix-turn-helix motif that contacts a pocket in the lumenal face of GCP6’s GRIP1 domain (Supplementary Figure S3H). This model agrees well with corresponding densities found in reconstructions of a recombinant human γ-TuRC from another study (Zimmermann et al., 2020), as well as the recombinant γ-TuRC determined here (Supplementary Figure S3I-J). Notably, GCP5’s insertion element loops back towards GCP5, suggesting that the remaining densities crossing towards GCP3 and GCP2 at positions 13-14 must constitute a separate polypeptide chain(s) (Zimmermann et al., 2020), although we were unable to discern its identity at the resolution we could achieve with our data.

The γ-TuRC also exhibits a MZT:GCP-NHD module-shaped density that associates with GCP3’s GRIP2 domain and is positioned right above γ-tubulin at position 1 at the so-called “seam” (Figure 1D). This feature may help “cap” the γ-TuRC during its assembly by blocking further GCP subunit additions past position 14 (Wieczorek et al., 2020b; a). Recently, this density has also been implicated in regulating nucleation by associating with α/β-tubulin in the early stages of tubulin assembly onto the γ-TuRC, acting as a “latch” that must be removed for complete γ-TuRC ring closure (Brito et al., 2024). Precisely which MZT:GCP-NHD module the latch density corresponds to is not entirely clear, although previous studies have tentatively assigned it as belonging to MZT1:GCP3-NHD (Würtz et al., 2022; Wieczorek et al., 2020a; Xu et al., 2024). However, given that there are only 5 copies of GCP3 in the γ-TuRC, our conclusion that the NEDD1 pinwheel contains 4 copies of MZT1:GCP3-NHD, together with the presence of a fifth MZT1:GCP3-NHD module in the lumenal bridge (Figure 1D), argue that the latch density must correspond to some other MZT:GCP-NHD module.

To reconcile this, we predicted structures of GCP3’s GRIP1 and GRIP2 domains and various MZT modules using AlphaFold. We found that a GCP3:MZT1:GCP5-NHD co-complex displayed good agreement with the seam density in our γ-TuRC (Figure 3B), as well as in previously-reported reconstructions (Würtz et al., 2022; Wieczorek et al., 2020a; Zimmermann et al., 2020). In contrast, the predicted GCP3:MZT1:GCP3-NHD model did not fit well into the seam density (Figure 3C). Further, the rigid body-fitted MZT1:GCP5-NHD model now satisfies a previously unexplained chemical cross-link between GCP5 residue K100 and position 1’s γ-tubulin residue K410 identified in the native human

γ-TuRC (Consolati et al., 2020) (Supplementary Figure S3D). Additionally, the flexible linker between GCP5’s GRIP1 domain and its NHD spans ∼80 residues (∼23 nm), satisfying the Euclidean distance between these two domains in the γ-TuRC (∼14 nm; Figure 3D-E). Our NEDD1-bound γ-TuRC structure therefore argues that the seam-associated “latch” MZT module belongs to MZT1:GCP5-NHD, and that GCP5 is involved in stabilizing long range interfaces across the γ-TuRC, analogous to how GCP6 incorporates into the lumenal bridge and uses the GCP6 belt feature to help stabilize GCP2 and GCP3 at the opposite side of the ∼30 nm-wide complex.

### The γ-TuRC can accommodate both a CDK5RAP2-containing CMG module and the NEDD1 pinwheel

Muroyama et al. proposed that cells can contain two populations of γ-TuRCs: those associated with NEDD1, which mainly function in microtubule anchoring, and those associated with the activating protein CDK5RAP2, which promotes microtubule nucleation from the complex (Muroyama et al., 2016). However, both NEDD1 and CDK5RAP2 are involved in the centrosomal localization of the γ-TuRC (Choi et al., 2010; Lüders et al., 2006). We note that the NEDD1 pinwheel does not obscure the interface between GRIP1 and GRIP2 domains in GCP2, corresponding to the known CDK5RAP2 binding sites, even at position 13 (Figure 1D) (Wieczorek et al., 2020b; a; Xu et al., 2024; Serna et al., 2024). It should therefore be possible for NEDD1 and CDK5RAP2 to simultaneously associate with the γ-TuRC, as hypothesized (Muroyama et al., 2016), but whether and how this might affect the γ-TuRC’s previously-reported conformational variability (Xu et al., 2024; Serna et al., 2024), are not yet clear.

To address these questions, we collected a large cryo-EM dataset of rec-γ-TuRC vitrified in the presence of residues 44-93 of CDK5RAP2, encompassing the so-called <u>γ-Tu</u>RC <u>n</u>ucleation <u>a</u>ctivating (γ-TuNA) motif (Supplementary Table S1) (Choi et al., 2010; Xu et al., 2024). Applying an analogous processing pipeline as with rec-γ-TuRC above increased the overall resolution of the CDK5RAP2-bound recombinant human γ-TuRC from 11 Å to 5.1 Å (Supplementary Figure S2C) (Xu et al., 2024). Gratifyingly, we also observed a pinwheel density in the new 3D reconstructions. Focused 3D classification led to a density map containing a better-resolved NEDD1 pinwheel, at the modest expense of resolution (6.9 Å; Figure 4A, Supplementary Figure S2C-D, G-H and Supplementary Table S2). In previous work, we showed that the CDK5RAP2-incubated recombinant γ-TuRC exhibits clear <u>C</u>DK5RAP2:<u>M</u>ZT2:<u>G</u>CP2-NHD (CMG) module formation, but only at complex position 13 (Xu et al., 2024). Our improved reconstructions confirmed the presence of the same CMG module even after subclassifying for NEDD1-containing densities (Figure 4A). Using this map, we built a molecular model of the recombinant human γ-TuRC simultaneously decorated by both CDK5RAP2 and NEDD1 (Figure 4A and Supplementary Table S3), which demonstrates that the NEDD1 pinwheel adopts a similar structure as in rec-γ-TuRC and, importantly, does not interfere with CMG module formation.

**Figure 4.**
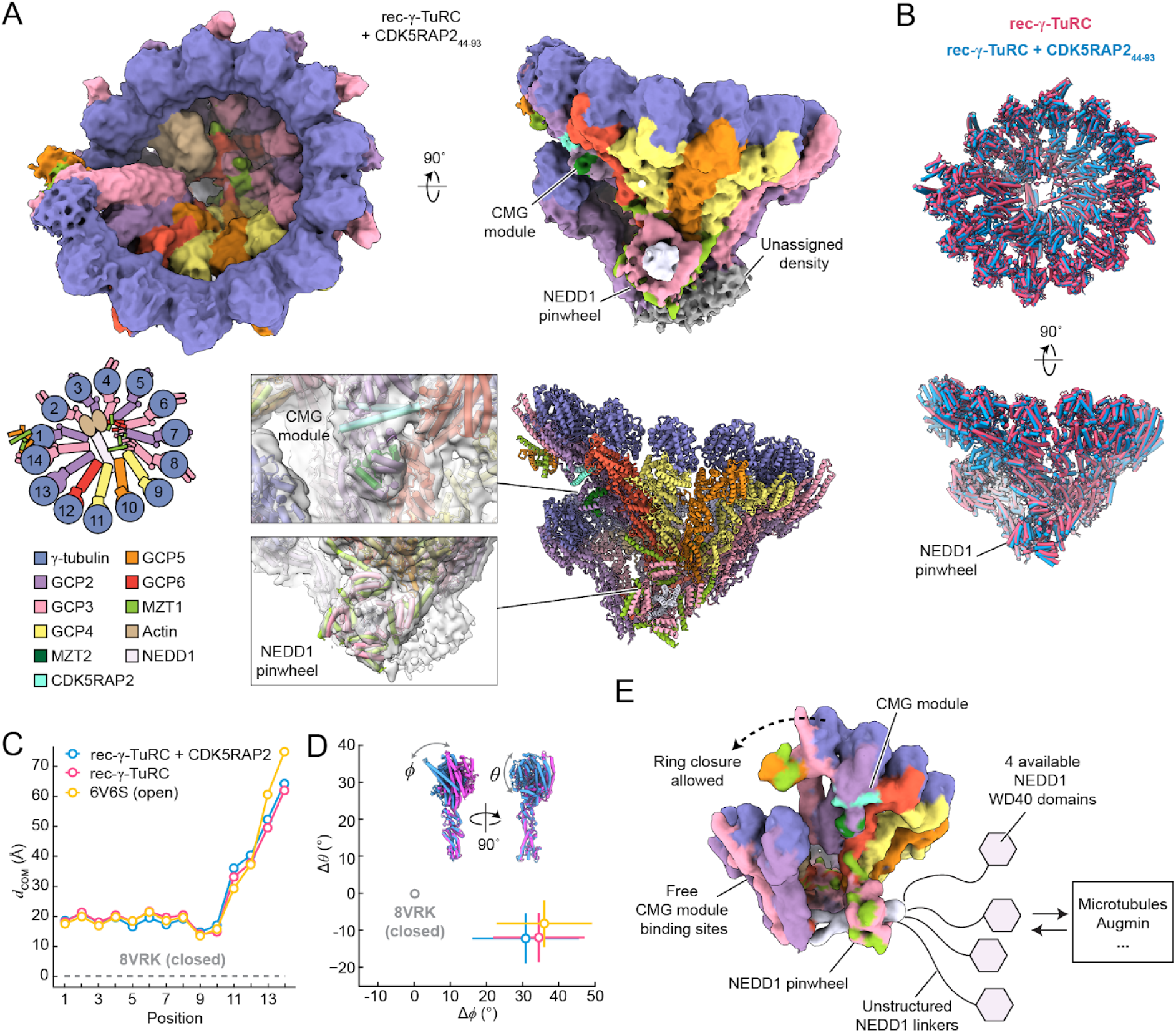
Both CDK5RAP2 and NEDD1 can simultaneously associate with the open γ-TuRC conformation. A) Top: two views of the consensus rec-γ-TuRC + CDK5RAP2 density map (surface representation). Unassigned, NEDD1 pinwheel and CMG module densities are indicated. Map resolution is 5.1 Å but is shown at a low threshold to include features with weaker density. Bottom left: A schematic top view of the γ-TuRC’s subunit organization. Bottom right: a refined molecular model of the rec-γ-TuRC + CDK5RAP2 (cartoon representation), with zoomed-in views for the CMG module and NEDD1 pinwheel in the density (transparent surface). B) Two cartoon representation views of superimposed rec-γ-TuRC (magenta) and rec-γ-TuRC + CDK5RAP2 (blue) models. The NEDD1 pinwheel is indicated (bottom). C) Plot of Euclidean center-of-mass distances (*d*_COM_) vs. γ-TuRC subunit position between the indicated γ-TuRC models (rec-γ-TuRC + CDK5RAP2, rec-γ-TuRC, and PDB: 6V6S as an “open” conformation control (Wieczorek et al., 2020b), all relative to PDB: 8VRK corresponding to the closed rec-γ-TuRC (Aher et al., 2024)). D) Plot of the average shift in θ vs. the shift in ϕ for helix H12 in γ-tubulins from each γ-TuRC described in (C), relative to γ-tubulins at the same positions in the closed γ-TuRC oligomer (grey circle, indicated). Standard errors in ϕ and θ are displayed as lines. The axes in (D) are scaled equally. Coloring in (D) follows the legend in (C). E) Model summarizing the findings in this study. The rec-γ-TuRC + CDK5RAP2 model has been converted to a 15 A low pass-filtered map and coloured accordingly. Unresolved WD40 domains stemming from the NEDD1 pinwheel and available to interact with binding partners are shown as hexagons. Free CMG module binding sites that should still be able to induce partial or full γ-tubulin ring closure are indicated.

### NEDD1 does not significantly alter the conformation of the γ-TuRC

What are the consequences of NEDD1 binding to the overall conformation of the γ-TuRC? Comparing rec-γ-TuRC models in the presence and absence of CDK5RAP2 did not reveal major differences in γ-TuRC subunit organization between the two NEDD1-bound complexes (Figure 4B). Notably, both models, which were built into classified maps containing strong NEDD1 pinwheel densities, are in the “open” γ-TuRC conformation, as judged from γ-tubulin:γ-tubulin Euclidean distances as well as GRIP2 domain rotation angles (Figure 4C-D), relative to the microtubule end-capped rec-γ-TuRC, which is instead in a “closed” conformation that matches the microtubule lattice (Aher et al., 2024). In sum, our results reveal that, while CDK5RAP2 and NEDD1 can simultaneously bind to the γ-TuRC, both rec-γ-TuRC and rec-γ-TuRC + CDK5RAP2 complexes are in the open conformation, suggesting that binding the NEDD1 pinwheel does not induce conformational changes in the γ-TuRC.

## Discussion

In this study, we determined the cryo-EM structure of NEDD1 bound to a fully-reconstituted human γ-TuRC. Our results demonstrate that NEDD1’s C-terminus adopts a tetrameric helical bundle that forms a critical interface with the GRIP1 domains of GCP2 and GCP3 next to the γ-TuRC seam. In turn, MZT1:GCP3-NHD subcomplexes encircle the NEDD1 tetramer in a pinwheel-shaped arrangement that docks onto the underside of the asymmetrically-positioned subunits GCP4, GCP5, and GCP6 in the cone-shaped complex. Our cryo-EM reconstructions also allowed us to assign several new densities in the γ-TuRC, including 1) an extension of the GCP6 belt that interacts with the fish tail of the NEDD1 pinwheel; 2) an insertion in GCP5’s GRIP1 domain that binds to the lumenal face of GCP6; and 3) a MZT1:GCP5-NHD module at the γ-TuRC seam.

Despite using full-length NEDD1 in our γ-TuRC reconstitution, only its C-terminal α-helices are resolved in the resulting cryo-EM density maps. The orientation of the NEDD1 helices in the pinwheel axle position NEDD1’s N-terminal unstructured and WD40 repeat domains away from the γ-TuRC core, freeing them to interact with microtubules and possibly other factors, such as the Augmin complex, which plays a key role in branching nucleation (Figure 4E) (Uehara et al., 2009; Zhang et al., 2022). Our observation that four NEDD1 molecules associate with each γ-TuRC may also help explain why oligomeric clusters of Augmin complexes are observed in single reconstituted microtubule branch sites (Zhang et al., 2022).

Furthermore, interactions between the γ-TuRC and Augmin are regulated via NEDD1 phosphorylation (Zhang et al., 2022). With the exception of S637, which is located next to the “fish tail” (Figure 2F), the main identified NEDD1 phosphorylation sites (S377, S405, S411, and a region encompassed by S557–S574 (Lüders et al., 2006; Pinyol et al., 2013; Sdelci et al., 2012; Gomez-Ferreria et al., 2012)) are not resolved in our structures. Rather, they all lie within the unstructured tether connecting the NEDD1 pinwheel with its WD40 domains, which also contains a putative Augmin binding site (Zhang et al., 2022). Although structural details regarding the γ-TuRC-Augmin interaction remain to be determined, our model of the NEDD1 pinwheel suggests that the region surrounding the γ-TuRC seam could constitute a potential binding site for microtubules and/or Augmin at the microtubule branch point.

We found that the presence of the NEDD1 pinwheel subcomplex does not influence the "open" conformation of the γ-TuRC. Instead, the large buried surface area provided by the NEDD1 pinwheel supports the hypothesis that NEDD1 acts as a mechanical anchor for the γ-TuRC that could also help orient the nucleated microtubule (Muroyama et al., 2016). That the NEDD1 pinwheel also does not prevent CDK5RAP2 from forming CMG modules on the γ-TuRC - at least at position 13 - provides a structural explanation for how NEDD1 and CDK5RAP2, both critical γ-TuRC attachment factors, can work in parallel to properly control γ-TuRC localization and activation at microtubule organizing centers (Muroyama et al., 2016; Fong et al., 2008). We note that, although we were not able to resolve CDK5RAP2-containing CMG modules at positions 3, 5, or 7 of the complex, as recently observed in other γ-TuRCs (Xu et al., 2024; Serna et al., 2024), the NEDD1 pinwheel associates only with the bottom of GCP4/5/6 GRIP1 domains on the opposite side of the complex. Accordingly, when performing our γ-TuRC conformation analysis (Figure 4C-D), we did not observe any obvious clashes between the NEDD1 pinwheel in the aligned rec-γ-TuRC and the relevant GRIP1 domains in the closed rec-γ-TuRC structure. Thus, our work argues that NEDD1 pinwheel binding to the complex should still allow γ-tubulin and the GRIP2 domains to undergo structural transitions into the partially-closed, fully CMG-decorated γ-TuRC (Xu et al., 2024; Serna et al., 2024), as well as to more closed microtubule end-capped γ-TuRC configurations (Brito et al., 2024; Aher et al., 2024) (Figure 4E).

## Acknowledgements

We thank Daniel Böhringer and Pavel Afanasyev from the Cryo-EM Knowledge Hub (ETH Zürich) for cryo-EM image processing advice, as well as Miroslav Peterek and Bilal Qureshi from ScopeM (ETH Zürich) for cryo-EM sample preparation and data collection training and support. Part of the cryo-EM data in this work was collected at the Simons Electron Microscopy Center (SEMC) at the New York Structural Biology Center, with support from the Simons Foundation (SF349247). We thank Linas Urnavicius and Roscislaw Krutyholowa for helpful discussions, as well as Christian Landolt and Alex Myczko (ETH Zürich) for IT support. This study includes calculations performed on the Euler cluster of ETH Zürich. Proteomics experiments were performed at the Functional Genomics Center Zurich (FGCZ) of University of Zürich and ETH Zürich. This work was supported by startup funds from the ETH Zürich, an SNSF Project Grant (#310030_208120), and an SNSF Starting Grant (#TMSGI3_211309), awarded to MW. TMK is grateful to the NIH (GM130234) for funding.

## Author Contributions

HMH, YX, TK, and MW designed the study. YX purified proteins used. HMH and YX collected new cryo-EM data. HMH, DZ, AX, AA, EW, and MW performed cryo-EM processing and model building. APC performed pulldown experiments. HMH, YX, APC and MW wrote the paper with contributions from all other authors. MW supervised the study and obtained funding.

## Materials and Methods

### Purification of recombinant human γ-TuRC (rec-γ-TuRC)

Polycistronic donor plasmid coding for human γ-tubulin, GCP2 and GCP3 (pACEBac1-γ-TuSC; Addgene plasmid # 178079; http://n2t.net/addgene:178079; RRID:Addgene_178079) and human γ-tubulin, GCP2, GCP3, GCP4, GCP5, GCP6, MZT1, ZZ-TEV-MZT2-3C-mEGFP, and actin (pACEBac1-γ-TuRC-GFP; Addgene plasmid # 178074; http://n2t.net/addgene:178074; RRID:Addgene_178074) were transformed into DH10MultiBacTurbo cells (ATG:biosynthetics GmbH) and transposition-positive colonies were selected and used to generate recombinant bacmids. Bacmids were transfected into Sf9 cells (Novagen) following the Bac-to-Bac manual (Invitrogen), baculoviruses were amplified twice, and fresh virus from γ-TuRC-GFP and γ-TuSC bacmids were mixed together at a 1:1 ratio. This virus mixture was used to infect 2 liters of High Five cells (Thermo Fisher Scientific) at a cell density of 3 × 10^6^ per ml for 60 h at 27°C. Cells were harvested by centrifugation at 1000 g, resuspended in 60 ml ice-cold lysis buffer (40 mM HEPES, pH 7.5, 150 mM KCl, 1 mM MgCl_2_, 10% glycerol (v/v), 0.1% Tween-20, 0.1 mM ATP, 0.1 mM GTP, 1 mM 2-mercaptoethanol, one cOmplete EDTA-free Protease Inhibitor Cocktail tablet (Roche), and 2 mM PMSF), and lysed by dounce homogenization on ice. The lysate was clarified at 322,000 g for 1 h at 4°C, 0.22-µm syringe–filtered, and loaded onto a 1 mL NHStrap column (Cytiva) previously coupled to 10 mg rabbit IgG (Innovative Biosciences; IRBIGGAP500MG) following the manufacturer’s instructions. The IgG column was washed with lysis buffer followed by gel filtration buffer (40 mM HEPES, pH 7.5, 150 mM KCl, 1 mM MgCl_2_, 10% glycerol (v/v), 0.1 mM GTP, and 1 mM 2-mercaptoethanol). An expression vector for TEV protease, pRK793, was a gift from David Waugh (Addgene plasmid 8827; http://n2t.net/addgene:8827; Research Resource Identifier: Addgene_8827; Kapust et al., 2001). TEV was expressed in BL21-CodonPlus (DE3)-RIL and purified using Ni-NTA and gel filtration following the methods described in (Ti et al., 2020). 1 mg of TEV protease (stored in 40 mM HEPES, pH 7.5, 30% (w/v) glycerol, 150 mM KCl, 1 mM MgCl_2_, and 3 mM 2-mercaptoethanol) was diluted into 1 ml of gel filtration buffer and injected onto the IgG column, and proteolysis was allowed to proceed for 2 h at 4°C. The digested eluate was pooled, concentrated with a 100 kDa cutoff spin filter (Millipore), and gel filtered over a Superose 6 Increase 10/300 GL column (Cytiva) pre-equilibrated in gel filtration buffer. Peak fractions were pooled and loaded onto two 2-ml sucrose gradient composed of 10%, 20%, 30%, and 40% sucrose (w/v) in gradient buffer (40 mM HEPES, pH 7.5, 150 mM KCl, 1 mM MgCl_2_, 0.01% Tween-20 (v/v), 0.1 mM GTP, and 1 mM 2-mercaptoethanol). The gradient was centrifuged at 50,000 rpm in a TLS-55 rotor at 4°C for 3 h with minimum acceleration and no break. Fractions were manually collected with a cut-off P1000 pipette tip and analyzed by SDS-PAGE followed by Coomassie staining and/or negative-stain TEM. Peak fractions were aliquoted, snap-frozen, and stored in liquid N2. Gradients were fractionated into 250 µl and analyzed by SDS-PAGE followed by Coomassie staining (Xu et al., 2024).

### Purification of recombinant, untagged CDK5RAP2 fragments

A bacterial expression construct for amino acids 44-93 of CDK5RAP2 lacking a GFP tag was previously described (Xu et al., 2024). The plasmid was transformed into BL21(DE3) pRIL *E. coli* cells (Stratagene) and His6-SUMO-TEV-CDK5RAP2_44-93_ expression was induced with 0.5 mM IPTG for 16 hours at 18°C. Cell pellets from 3 L culture were resuspended in 45 mL Ni-NTA lysis buffer (50 mM sodium phosphate, pH 8.0, 300 mM NaCl, 15 mM imidazole, 0.1% (v/v) Tween-20 and 1 mM 2-mercaptoethanol) and lysed by three passes through an Emulsiflex C-5 (Avestin). Lysate was clarified at 35,000 rpm in a Type 45 Ti rotor (Beckman) for 45 min at 4°C and the supernatant was mixed with 3 mL Ni-NTA agarose (Qiagen) pre-equilibrated in lysis buffer. The resin was then washed extensively with lysis buffer. Protein was eluted with Ni-NTA elution buffer (lysis buffer containing an additional 485 mM imidazole). Peak fractions were identified by Bradford assay, pooled, concentrated to ∼2.5 mL with a 3 kDa cutoff spin filter (Millipore), and applied to PD-10 desalting column (Cytiva) pre-equilibrated with lysis buffer to remove imidazole. The elution was incubated with 1 mg TEV protease for 2 hr at 4°C, and injected into a 1 mL HisTrap column (Cytiva) pre-equilibrated with lysis buffer. Peak fractions were identified by Bradford assay, pooled, concentrated to ∼ 1 mL at ∼ 50 mg/mL with the 3 kDa cutoff spin filter (Millipore). Untagged CDK5RAP2_44-93_ was then further purified over a HiLoad 16/600 Superdex 75 column pre-equilibrated in gel filtration buffer (40 mM HEPES, pH 7.5, 150 mM NaCl, 1 mM MgCl_2_ and 2 mM 2-mercaptoethanol). Peak fractions were identified by SDS-PAGE, pooled, concentrated, supplemented with sucrose to 10% (w/v), aliquoted, snap frozen in liquid nitrogen, and stored at –80°C. After TEV cleavage, the resulting CDK5RAP2_44-93_ peptide contains an N-terminal SNIGSGGTPSTGSSLPNVSEE sequence as a remnant of the linking sequence between the TEV cleavage site and the multiple cloning site in the modified expression vector. LC-MS/MS was used to confirm the identity of purified CDK5RAP2_44-93_, as well as a lack of phosphorylated peptides, as expected for a bacterially-expressed protein (Xu et al., 2024).

### Cryo-EM sample preparation

EM grids were prepared with Ultrathin Carbon Film on a Lacey Carbon Support Film (400 mesh, Copper Ted Pella, Inc) or by overlaying Quantifoil 300 mesh (copper, R 2/2) holy carbon grids with homemade continuous carbon film. Lacey grids were glow discharged at 30 mA for 45 seconds, and at 15 mA for 15 seconds for Quantifoil grids prepared with homemade carbon support films. Following, the grids were transferred using metal forceps onto an ice-cold block. rec-γ-TuRC samples were pre-incubated with 2 μM CDK5RAP2_44-93_ for 10 minutes on ice prior to grid application. A 2.5 μL aliquot of mixed sample was applied to the grid for 5 minutes, followed by manual blotting. This process was repeated 8 times. After the final application, the grid was washed by incubating twice with 20 μL of washing buffer (40 mM HEPES, pH 7.5, 150 mM KCl, 1 mM MgCl_2_, 1 mM 2-mercaptoethanol, 0.01% Tween-20, 0.1 mM GTP) for 1 min. An additional 3.5 μL of washing buffer was applied onto the grid. The grid was transferred to a Vitrobot Mark IV (ThermoFisher Scientific), blotted for 4 seconds with a blot force of -1 at 100% humidity and 4°C, plunge-frozen in a liquid ethane/propane mixture, and stored in liquid nitrogen until screening in the ScopeM facility at ETH (Zürich, Switzerland).

### Cryo-EM data collection

For rec-γ-TuRC + CDK5RAP2, 192,360 new movies were collected on a TFS Titan Krios G3i (2) FEG operated with a Gatan K3 in CDS mode and a slit width of 20 eV on a GIF BioQuantum energy filter. These datasets were combined with four previously-described datasets for analysis (Xu et al., 2024). Full cryo-EM data collection statistics are listed in Supplementary Table S2. Automatic data collection was performed with “Faster acquisition mode” in EPU software (Thermo Fisher Scientific).

### Cryo-EM image processing

#### rec-γ-TuRC

Particles were first autopicked in RELION 4.0 from micrographs in (Aher et al., 2024) using 2D classes generated from a small subset of manually-picked γ-TuRC particles as picking templates. Autopicked particles were then subjected to multiple rounds of 2D and 3D classification to form a roughly cleaned stack of 1,990,604 γ-TuRC particles. The particles were imported into CryoSPARC and classified by heterogeneous refinement using two copies of γ-TuRC averaged from the particle stack and five “noise” density maps generated by premature cancellation of a 90-particle *ab initio* refinement as references. This was performed in multiple rounds until minimal proportions of particles were being assigned to the noise classes, revealing a subset of 584,330 particles. A final heterogeneous refinement was performed using three copies of an average of the particle subset, as well as three copies of the average of the whole dataset, revealing a subset of 266,675 particles with better quality γ-TuRC density. The final particles were subjected to local refinement using a γ-TuRC-shaped mask to generate the final consensus rec-γ-TuRC reconstruction at 4.7 Å resolution (Supplementary Figure S2A-B & E-F).

We also tested different software tools to analyze the heterogeneity and occupancy of the NEDD1 pinwheel, including those developed in RELION-5 (Blush regularization, DynaMight, and MultiBody refinements (Burt et al., 2024)), cryoDRGN (Zhong et al., 2021), and OccuPy (Forsberg et al., 2023), none of which significantly improved tertiary or secondary structure features within the pinwheel density.

#### rec-γ-TuRC + CDK5RAP2

6,000 movies from each dataset were first used to estimate a gain reference for each rec-γ-TuRC + CDK5RAP2 dataset using “relion_estimate_gain” (Burt et al., 2024). Movies were then imported into CryoSPARC for patch motion correction and patch contrast transfer function (CTF) estimation (Punjani et al., 2017). A blob picker with an elliptical blob size of 280-380 Å was used to obtain a curated set of particles, selected through 2D classification and visual inspection of micrographs. Exposures were chosen based on CTF fit resolution (2-18 Å), astigmatism (0-2000 Å), and relative ice thickness (0-1.8).

A ResNet16 neural network was trained on the curated particle set extracted from 16x downsampled micrographs with TOPAZ (Bepler et al., 2020). TOPAZ models were generated for each independent dataset, and the software was used to pick and extract particles using a box size of 196. The TOPAZ-picked particles were subjected to 2D classification in CryoSPARC, and particles were selected based on 2D classes displaying the distinctive γ-TuRC shape.

To increase particle numbers, TOPAZ training and picking were repeated using the selected particles. Selected particles were extracted using a box size of 848 pixels, binned to 144 or 256 pixels, and subjected to 2D classification. Any repeating particle coordinates within 200 Å of each other were removed using the “remove duplicates” feature in CryoSPARC. The resulting 296,981 particles were then subjected to *ab initio* reference generation expecting three classes. However, only one γ-TuRC-like class resulted from this run; the other two classes converged as noise references. All particles were then subjected to heterogeneous refinement with “force hard classification” turned on and using the *ab inito* maps as starting references. The particles classified into the γ-TuRC-like class were re-extracted at a box size of 384 pixels and subjected to non-uniform refinement.

To generate a consensus reconstruction of rec-γ-TuRC + CDK5RAP2 with improved NEDD1 density, 3D classification was performed in CryoSPARC, with a mask focusing on the pinwheel-shaped density. This classification, using two classes and filtered at 12 Å, aimed to uncover rare or hidden conformations. The first class contained 77,282 particles, while the second had 71,778 particles. Both classes were independently refined via local refinement, producing a final consensus map of the rec-γ-TuRC + CDK5RAP2 complex with well-resolved NEDD1 pinwheel density from the second class (Supplementary Figure S2C-D & G-H).

### Model building

For the rec-γ-TuRC model, a combination of structure predictions were first generated using AlphaFold 3 (Abramson et al., 2024). For instance, NEDD1 (Uniprot Accession: Q8NHV4), MZT1 (Uniprot Accession: Q08AG7), and GCP3-NHD (Uniprot Accession: Q96CW5) protein sequences were used to assemble the pinwheel (Figure 1D). GCP2 (Uniprot Accession: Q9BSJ2), GCP3 (Uniprot Accession: Q96CW5), GCP4 (Uniprot Accession: Q9UGJ1), GCP5 (Uniprot Accession: Q96RT8), GCP6 (Uniprot Accession: B2RWN4), and MZT1 (Uniprot Accession: Q08AG7), together with γ-tubulin (Uniprot Accession: P23258) :GCP3:MZT1:GCP5-NHD and γ-tubulin:GCP3:MZT1:GCP3-NHD, were all used to interpret the consensus map. Lumenal bridge components were similarly predicted with AlphaFold 3.

The atomic model of the native human γ-TuRC (PDB ID: 6V6S) was fitted into the rec-γ-TuRC consensus map (Figure 1A). The different AlphaFold predictions were initially aligned to this model using the “matchmaker” tool in ChimeraX and then manually corrected in Coot. Due to insufficient resolution for model building, side chains were trimmed to β-carbons, and the resulting model was real-space refined in PHENIX (Afonine et al., 2018).

For the rec-γ-TuRC + CDK5RAP2 model, a starting AlphaFold 3 structure containing GCP2, GCP6, MZT2 (Q6NZ67), and two copies of residues 44-93 of CDK5RAP2 (I3LKY1) was docked into the rec-γ-TuRC + CDK5RAP2 map using the “fit in map” function of UCSF ChimeraX (Goddard et al., 2018). All other subunits originated from the rec-γ-TuRC model above. Model domains were positioned into corresponding local density using the “Fit all chains to Map” tool in Coot, and any loops or structural features not supported by local density were removed. Due to insufficient resolution for model building, side chains were trimmed to β-carbons, and the resulting model was real-space refined in PHENIX (Afonine et al., 2018).

Refinement statistics for rec-γ-TuRC and rec-γ-TuRC + CDK5RAP2 models are available in Supplementary Table S3.

### NEDD1 pull-down experiments

Mammalian expression pcDNA3.1 vectors encoding wild-type, E598A/D602A/E605A and F603A/F622A NEDD1 with N-terminal Myc-SBP (streptavidin-binding peptide) tag were synthetized via gene synthesis (Genewiz). The canonical NEDD1 protein isoform Q8NHV4-1 was used. 2.5 x 10^6^ HEK293T cells, cultured in DMEM containing 10% FBS and 1X penicillin/streptomycin, were seeded in 10 cm plates and transfected 24 h later using PEI Max (1:3 of DNA-to-PEI mass ratio). Cells were transfected with 20 µg of total plasmid and 24 h after transfection the medium was changed. Another 24 h later, cells were harvested and lysed for 30 min on ice with lysis buffer (50 mM Tris-HCl pH 7.5, 150 mM NaCl, 0.1% Triton X-100, 10% Glycerol, 2 mM DTT) supplemented with EDTA-free protease inhibitor (Roche), 1 mM sodium fluoride, 1 mM ꞵ-glycerophosphate, and 1 mM sodium pyrophosphate. After shearing the lysates with a syringe and a needle (collecting the lysate for 6 times on ice), crude lysates were centrifuged at 16,000 g for 15 min at 4 C and their total protein amount normalized with the Bradford protein assay. The cleared lysate was rotated for 1 h at 4°C in presence of 50 mL of Streptavidin Sepharose High Performance (Cytiva). Beads were washed three times with lysis buffer. Bound proteins were eluted with 50 mL 2 xSDS protein sample buffer. For further analysis, proteins were separated by SDS-PAGE and detected by Western blotting. c-Myc monoclonal antibody (clone 9E10, Cat# MA1-980, Invitrogen) and anti-γ-tubulin monoclonal antibody (clone GTU-88, Cat# T5326, Sigma-Aldrich) were used at 1:1000 and 1:2000 dilutions respectively. HRP-conjugated goat anti-mouse IgG secondary antibody (Cat#31430, Invitrogen) was used at 1:5000. Untransfected cells served as negative control.

### Helical parameter analysis

Models for rec-γ-TuRC, rec-γ-TuRC + CDK5RAP2, and the human native γ-TuRC in the open conformation (PDB 6V6S; (Wieczorek et al., 2020b)) were aligned to a structure of γ-TuRC bound to the microtubule end (PDB 8RVK; (Aher et al., 2024)) via the GRIP1 domains of GCP3 at positions 2, 4, 6, and 8 using the align command in PyMoL. The aligned structures were subsequently read out into PDB format and analyzed using a set of previously-described custom python scripts that can be found at: https://github.com/wieczoreklab/gTuRC-YX/blob/main/gTuRC-geometry. Specifically, the aligned PDB files were loaded and coordinates of all Cα atoms in the structures were extracted using biophython (Cock et al., 2009). The position of each γ-tubulin was given by the center of mass (CoM), i.e., the mean of its Cα atoms. The CoM was used to analyze structural changes due to its robustness and stability concerning conformational changes of individual γ-tubulins. The Euclidean distance between the CoM of the reference rec-γ-TuRC in the closed conformation and all other γ-TuRC structures at each position was calculated to analyze changes between complexes.

To examine changes in orientation of individual γ-tubulins across the structures, the CoM of the Cα’s of the first half and second half of γ-tubulin helix 12 was used to define a vector describing tilt and rotation of each γ-tubulin. The coordinates were transformed to spherical coordinates, where phi (ϕ) denotes the angle of the vectors projected in x-y-plane, while theta (Ө) describes the tilt away from the z-axis (the length of the vectors are set to 1 for simplicity). Normalization of theta and phi was done by subtraction, resulting in the direction of the change in orientation of each γ-tubulin with respect to the closed rec-γ-TuRC reference structure.

---

**Figure S1.**
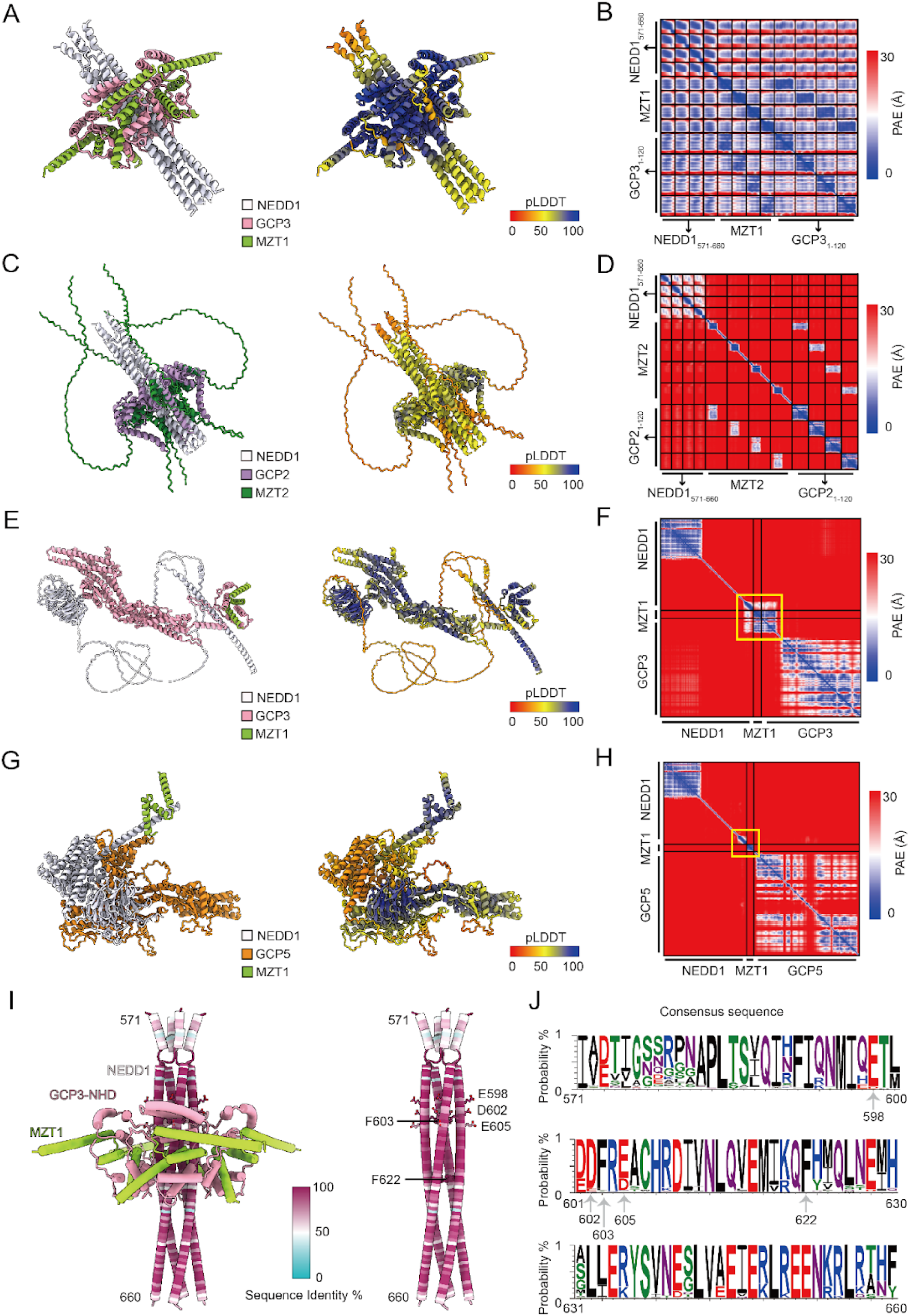
AlphaFold predictions and NEDD1 conservation analysis. A) Cartoon representation AlphaFold 3 prediction of four copies each of human NEDD1(571-660), MZT1, and GCP3(1-120) coloured by subunit (left) and pLDDT (right). B) Partial alignment error plot for the prediction in A). C) Cartoon representation AlphaFold 3 prediction of four copies each of human NEDD1(571-660), MZT2A, and GCP2(1-120) coloured by subunit (left) and pLDDT (right). D) Partial alignment error plot for the prediction in C). E) Cartoon representation AlphaFold 3 prediction of full-length *A. thaliana* NEDD1, MZT1A, and GCP3 coloured by subunit (left) and pLDDT (right). F) Partial alignment error plot for the prediction in E). G) Cartoon representation AlphaFold 3 prediction of full-length *A. thaliana* NEDD1, MZT1A, and GCP5A coloured by subunit (left) and pLDDT (right). H) Partial alignment error plot for the prediction in G). Yellow boxes in F) and H) highlight expected regions for formation of NEDD1:MZT:GCP-NHD subcomplexes. I) Cartoon representation of AlphaFold 3 model of the NEDD1 pinwheel in A). MZT1 and GCP3-NHD are coloured according to the legend in A); NEDD1 is coloured according to conservation as scored in the colour key. A multiple sequence alignment of 141 annotated protein sequences across various species was used to score conservation. Residues mutated in this study are shown in stick representation on the right-hand model of NEDD1 alone. J) Conservation of NEDD1 residues 571-660. Generated using WebLogo (Crooks et al., 2004).

**Figure S2.**
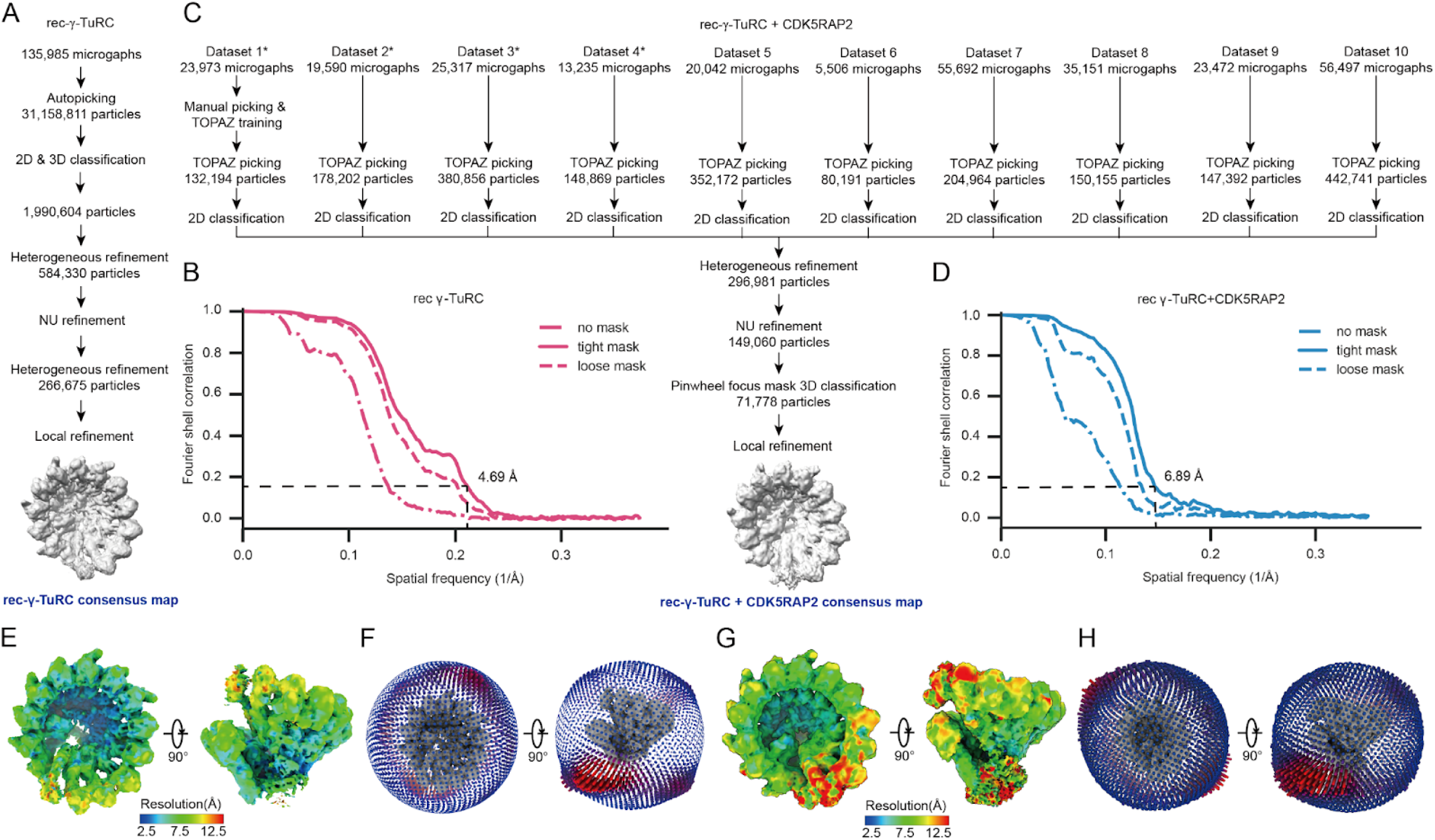
Cryo-EM processing pipeline. A) Summary of the new rec-γ-TuRC cryo-EM data processing strategy. Cryo-EM collection details were reported in (Aher et al., 2024). B) Gold-standard Fourier shell correlation (FSC) plot for the consensus rec-γ-TuRC density map. The FSC at 0.143 is indicated by a dashed line. C) Summary of new rec-γ-TuRC + CDK5RAP2 cryo-EM data processing strategy. Cryo-EM collection details for Datasets 1-4 (marked by asterisks) were reported in (Xu et al., 2024). D) Gold-standard Fourier shell correlation (FSC) plot for the consensus rec-γ-TuRC + CDK5RAP2 density map. The FSC at 0.143 is indicated by a dashed line. E) Two views of the consensus rec-γ-TuRC density map analyzed by CryoSPARC, showing a resolution distribution ranging from 2.5 to >12.5 Å. F) Two views of the particle angular distribution overlaid onto the rec-γ-TuRC consensus map. G) Two views of the consensus rec-γ-TuRC + CDK5RAP2 density map analyzed by CryoSPARC, showing a resolution distribution ranging from 2.5 to >12.5 Å. H) Two views of the particle angular distribution overlaid onto the rec-γ-TuRC + CDK5RAP2 consensus map.

**Figure S3.**
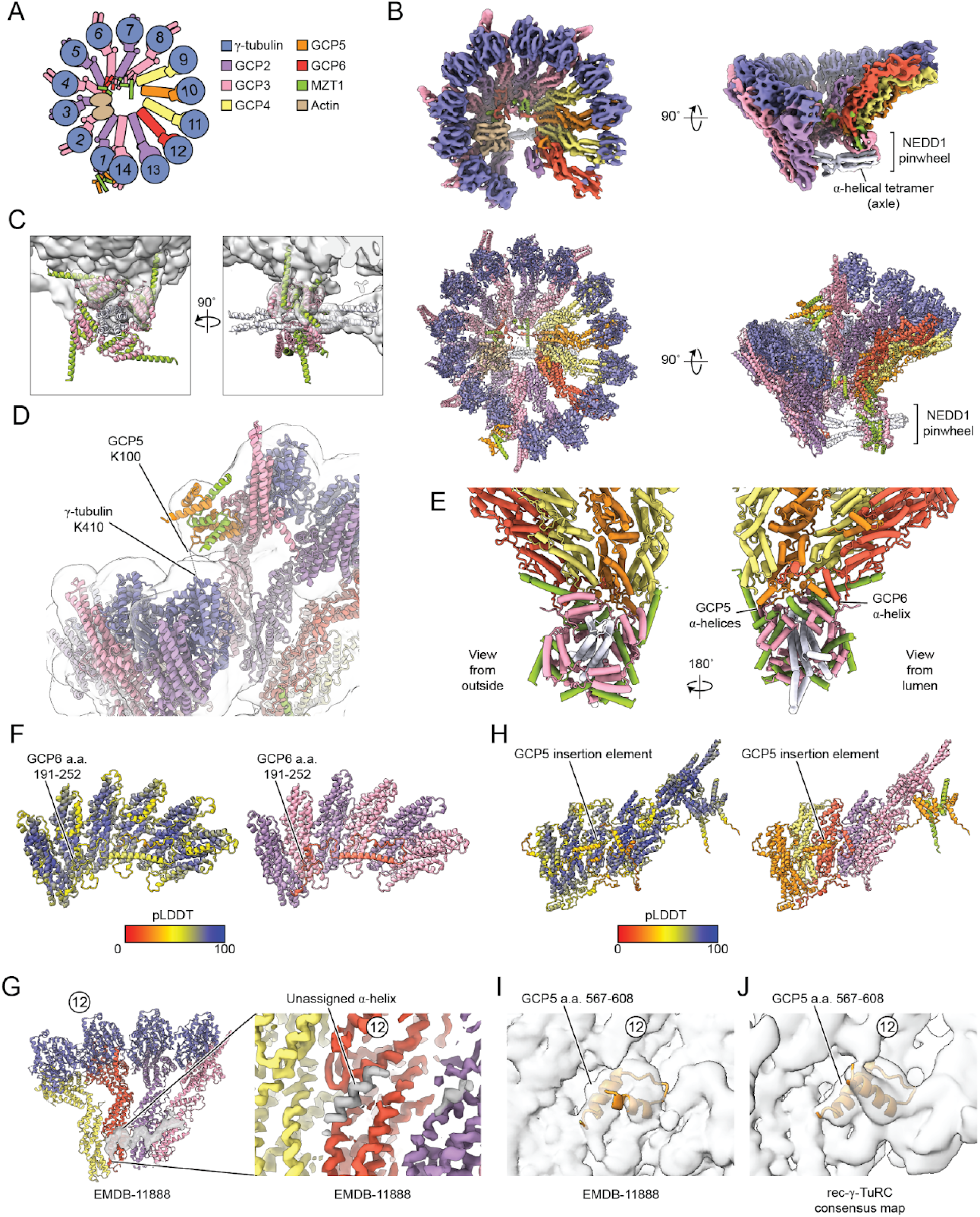
Details regarding rec-γ-TuRC consensus reconstruction and model building. A) Schematic of the γ-TuRC highlighting subunit composition and numbering across the complex. B) Top: two views of the rec-γ-TuRC consensus map showing higher resolution features. Map was sharpened in CryoSPARC and post-processed with EMready (He et al., 2023). Bottom: two views of the refined rec-γ-TuRC model including the NEDD1 pinwheel. C) Two views of the NEDD1 pinwheel predicted by AlphaFold 3 (cartoon representation) fitted into the pinwheel density in the rec-γ-TuRC consensus map (transparent surface). D) Cartoon representation view of MZT1:GCP5-NHD at the rec-γ-TuRC seam with the consensus density map in transparent surface representation. GCP5 K100 and γ-tubulin K410 identified as crosslinked residues in the native human γ-TuRC are indicated (Consolati et al., 2020). E) Two views of the NEDD1 pinwheel docked onto the underside of GCP4-6 at positions 9-12 of the γ-TuRC. GCP5-and GCP6-specific α-helices that contact the pinwheel are indicated. F) Cartoon representation AlphaFold 3 prediction of three copies each of GCP2 and GCP3 GRIP1 domains together with the GCP6 belt and residues 191-252, coloured by pLDDT (left) and subunit (right). G) Segmented surface representation of the previously-described helical element lining the lumenal face of GCP6, 2, and 3 in EMDB-11888 (Zimmermann et al., 2020). Map was post-processed with EMready (He et al., 2023). The γ-TuRC subunits from the same study are shown in cartoon representation for reference. A zoomed-in view of an unassigned helix contacting GCP6 is shown on the right and at a higher threshold. H) Cartoon representation AlphaFold 3 prediction of GRIP1 domains of GCP4, GCP5 (including NHD), GCP6 and GCP2, as well as MZT1 and the GRIP1 and GRIP2 domains of GCP3, coloured by pLDDT (left) and subunit (right). The GCP5 insertion element that contacts the lumenal face of GCP6 is indicated. I) The GCP5 insertion (a.a. 567-608) is shown in cartoon representation fitted into the EMDB-11888 density map (Zimmermann et al., 2020). J) The GCP5 insertion modeled in this study (a.a. 567-608) is shown in cartoon representation in the rec-γ-TuRC density map. γ-TuRC position 12 corresponding to GCP6 is indicated in panels G), I) and J) for reference.

**Table S1.** Cryo-EM data collection.

|  | rec-γ-TuRC + CDK5RAP2 |  |  |  |  |  |
| --- | --- | --- | --- | --- | --- | --- |
|  | Dataset 5 | Dataset 6 | Dataset 7 | Dataset 8 | Dataset 9 | Dataset 10 |
| Voltage (kV) | 300 | 300 | 300 | 300 | 300 | 300 |
| Magnification | 130,000X | 130,000X | 130,000X | 130,000X | 130,000X | 130,000X |
| Pixel size (Å/px) | 0.64 | 0.64 | 0.64 | 0.64 | 0.64 | 0.64 |
| Camera | K3 | K3 | K3 | K3 | K3 | K3 |
| GIF slit | 20 eV | 20 eV | 20 eV | 20 eV | 20 eV | 20 eV |
| Mode | CDS | CDS | CDS | CDS | CDS | CDS |
| Software | EPU | EPU | EPU | EPU | EPU | EPU |
| Spot size | 6 | 5 | 6 | 6 | 6 | 7 |
| Beam diameter (μm) | 0.82 | 0.78 | 0.81 | 0.86 | 0.80 | 0.57 |
| # of movies | 20,042 | 5,506 | 55,692 | 35,151 | 23,472 | 56,497 |
| # of frames | 50 | 50 | 50 | 46 | 46 | 40 |
| Exposure time (s) | 0.9 | 0.9 | 1 | 1 | 1 | 1 |
| Dose Rate (e<sup>-</sup>/px/s) | 31 | 27 | 24 | 23 | 25 | 26 |
| Total dose (e<sup>-</sup>/Å<sup>2</sup>) | 66 | 61 | 59 | 56 | 65 | 64 |
| Defocus range (μm) | -0.8/-2.4 | -0.8/-2.4 | -0.8/-2.4 | -0.8/-2.4 | -0.9/-2.4 | -1/-2.4 |

**Table S2.** Cryo-EM data processing statistics.

|  | **rec-γ-TuRC**<br>**consensus map** | **rec-γ-TuRC + CDK5RAP2**<br>**consensus map** |
| --- | --- | --- |
| Processing pixel size (Å/pixel) | 1.32 | 1.41 |
| Box size (pixels) | 368 | 384 |
| Symmetry imposed | C1 | C1 |
| Number of particles | 266,675 | 71,778 |
| Resolution (Å)<br>(FSC 0.143 cutoff) | 4.7 (CS) | 6.8 (CS) |
| Resolution range (Å) (ResMap) | (3-11) | NC |
| Model building | Yes | Yes |
| Accession number | EMDB-XXXXX | EMDB-XXXXX |

\*NC = Not calculated
\*CS = CryoSPARC

**Table S3.** Model building and refinement statistics.

|  | **rec-γ-TuRC** | **rec-γ-TuRC + CDK5RAP2** |
| --- | --- | --- |
| Cross-correlation | 0.43 | 0.38 |
| Number of chains | 45 | 48 |
| Number of residues | 17,575 | 17,794 |
| All atom clash score | 6.26 | 7.03 |
| Outliers (%) | 0.23 | 0.22 |
| Allowed (%) | 2.05 | 2.07 |
| Favoured (%) | 97.72 | 97.70 |
| Bond length (Å) | 0.002 | 0.002 |
| Bond angles (°) | 0.510 | 0.532 |
| Accession number | PDB: XXXX | PDB: XXXX |

## Notes

### Competing Interest Statement

The authors have declared no competing interest.

## References

Abramson, J., J. Adler, J. Dunger, R. Evans, T. Green, A. Pritzel, O. Ronneberger, L. Willmore, A.J. Ballard, J. Bambrick, S.W. Bodenstein, D.A. Evans, C.-C. Hung, M. O’Neill, D. Reiman, K. Tunyasuvunakool, Z. Wu, A. Žemgulytė, E. Arvaniti, C. Beattie, O. Bertolli, A. Bridgland, A. Cherepanov, M. Congreve, A.I. Cowen-Rivers, A. Cowie, M. Figurnov, F.B. Fuchs, H. Gladman, R. Jain, Y.A. Khan, C.M.R. Low, K. Perlin, A. Potapenko, P. Savy, S. Singh, A. Stecula, A. Thillaisundaram, C. Tong, S. Yakneen, E.D. Zhong, M. Zielinski, A. Žídek, V. Bapst, P. Kohli, M. Jaderberg, D. Hassabis, and J.M. Jumper. 2024. Accurate structure prediction of biomolecular interactions with AlphaFold 3. Nature. 630:493–500.

Afonine, P.V., B.K. Poon, R.J. Read, O.V. Sobolev, T.C. Terwilliger, A. Urzhumtsev, and P.D. Adams. 2018. Real-space refinement in PHENIX for cryo-EM and crystallography. Acta Crystallogr D Struct Biol. 74:531–544.

Aher, A., L. Urnavicius, A. Xue, K. Neselu, and T.M. Kapoor. 2024. Structure of the γ-tubulin ring complex-capped microtubule. Nat. Struct. Mol. Biol. doi:10.1038/s41594-024-01264-z.

Bepler, T., K. Kelley, A.J. Noble, and B. Berger. 2020. Topaz-Denoise: general deep denoising models for cryoEM and cryoET. Nat. Commun. 11:5208.

Brito, C., M. Serna, P. Guerra, O. Llorca, and T. Surrey. 2024. Transition of human γ-tubulin ring complex into a closed conformation during microtubule nucleation. Science. 383:870–876.

Burt, A., B. Toader, R. Warshamanage, A. von Kügelgen, E. Pyle, J. Zivanov, D. Kimanius, T.A.M. Bharat, and S.H.W. Scheres. 2024. An image processing pipeline for electron cryo-tomography in RELION-5. bioRxiv. 2024.04.26.591129. doi:10.1101/2024.04.26.591129.

Choi, Y.-K., P. Liu, S.K. Sze, C. Dai, and R.Z. Qi. 2010. CDK5RAP2 stimulates microtubule nucleation by the gamma-tubulin ring complex. J. Cell Biol. 191:1089–1095.

Cock, P.J.A., T. Antao, J.T. Chang, B.A. Chapman, C.J. Cox, A. Dalke, I. Friedberg, T. Hamelryck, F. Kauff, B. Wilczynski, and M.J.L. de Hoon. 2009. Biopython: freely available Python tools for computational molecular biology and bioinformatics. Bioinformatics. 25:1422–1423.

Consolati, T., J. Locke, J. Roostalu, Z.A. Chen, J. Gannon, J. Asthana, W.M. Lim, F. Martino, M.A. Cvetkovic, J. Rappsilber, A. Costa, and T. Surrey. 2020. Microtubule Nucleation Properties of Single Human γTuRCs Explained by Their Cryo-EM Structure. Dev. Cell. 53:603–617.e8.

Cota, R.R., N. Teixidó-Travesa, A. Ezquerra, S. Eibes, C. Lacasa, J. Roig, and J. Lüders. 2017. MZT1 regulates microtubule nucleation by linking γTuRC assembly to adapter-mediated targeting and activation. J. Cell Sci. 130:406–419.

Cunha-Ferreira, I., A. Chazeau, R.R. Buijs, R. Stucchi, L. Will, X. Pan, Y. Adolfs, C. van der Meer, J.C. Wolthuis, O.I. Kahn, P. Schätzle, M. Altelaar, R.J. Pasterkamp, L.C. Kapitein, and C.C. Hoogenraad. 2018. The HAUS complex is a key regulator of non-centrosomal microtubule organization during neuronal development. Cell Rep. 24:791–800.

Dendooven, T., S. Yatskevich, A. Burt, Z.A. Chen, D. Bellini, J. Rappsilber, J.V. Kilmartin, and D. Barford. 2024. Structure of the native γ-tubulin ring complex capping spindle microtubules. Nat. Struct. Mol. Biol. doi:10.1038/s41594-024-01281-y.

Fong, K.-W., Y.-K. Choi, J.B. Rattner, and R.Z. Qi. 2008. CDK5RAP2 is a pericentriolar protein that functions in centrosomal attachment of the gamma-tubulin ring complex. Mol. Biol. Cell. 19:115–125.

Forsberg, B.O., P.N.M. Shah, and A. Burt. 2023. A robust normalized local filter to estimate compositional heterogeneity directly from cryo-EM maps. Nat. Commun. 14:1–11.

Goddard, T.D., C.C. Huang, E.C. Meng, E.F. Pettersen, G.S. Couch, J.H. Morris, and T.E. Ferrin. 2018. UCSF ChimeraX: Meeting modern challenges in visualization and analysis. Protein Sci. 27:14–25.

Gomez-Ferreria, M.A., M. Bashkurov, A.O. Helbig, B. Larsen, T. Pawson, A.-C. Gingras, and L. Pelletier. 2012. Novel NEDD1 phosphorylation sites regulate γ-tubulin binding and mitotic spindle assembly. J. Cell Sci. 125:3745–3751.

Goshima, G., M. Mayer, N. Zhang, N. Stuurman, and R.D. Vale. 2008. Augmin: a protein complex required for centrosome-independent microtubule generation within the spindle. J. Cell Biol. 181:421–429.

Haren, L., M.-H. Remy, I. Bazin, I. Callebaut, M. Wright, and A. Merdes. 2006. NEDD1-dependent recruitment of the gamma-tubulin ring complex to the centrosome is necessary for centriole duplication and spindle assembly. J. Cell Biol. 172:505–515.

He, J., T. Li, and S.-Y. Huang. 2023. Improvement of cryo-EM maps by simultaneous local and non-local deep learning. Nat. Commun. 14:3217.

Hotta, T., Z. Kong, C.-M.K. Ho, C.J.T. Zeng, T. Horio, S. Fong, T. Vuong, Y.-R.J. Lee, and B. Liu. 2012. Characterization of the Arabidopsis augmin complex uncovers its critical function in the assembly of the acentrosomal spindle and phragmoplast microtubule arrays. Plant Cell. 24:1494–1509.

Huang, T.-L., H.-J. Wang, Y.-C. Chang, S.-W. Wang, and K.-C. Hsia. 2020. Promiscuous binding of microprotein Mozart1 to γ-tubulin complex mediates specific subcellular targeting to control microtubule array formation. Cell Rep. 31:107836.

Kimanius, D., L. Dong, G. Sharov, T. Nakane, and S.H.W. Scheres. 2021. New tools for automated cryo-EM single-particle analysis in RELION-4.0. Biochem. J. 478:4169–4185.

Kollman, J.M., A. Merdes, L. Mourey, and D.A. Agard. 2011. Microtubule nucleation by γ-tubulin complexes. Nat. Rev. Mol. Cell Biol. 12:709–721.

Liu, L., and C. Wiese. 2008. Xenopus NEDD1 is required for microtubule organization in Xenopus egg extracts. J. Cell Sci. 121:578–589.

Liu, P., E. Zupa, A. Neuner, A. Böhler, J. Loerke, D. Flemming, T. Ruppert, T. Rudack, C. Peter, C. Spahn, O.J. Gruss, S. Pfeffer, and E. Schiebel. 2019. Insights into the assembly and activation of the microtubule nucleator γ-TuRC. Nature. 578:467–471.

Liu, T., J. Tian, G. Wang, Y. Yu, C. Wang, Y. Ma, X. Zhang, G. Xia, B. Liu, and Z. Kong. 2014. Augmin Triggers Microtubule-Dependent Microtubule Nucleation in Interphase Plant Cells. Curr. Biol. 24:2708–2713.

Lüders, J., U.K. Patel, and T. Stearns. 2006. GCP-WD is a gamma-tubulin targeting factor required for centrosomal and chromatin-mediated microtubule nucleation. Nat. Cell Biol. 8:137–147.

Manning, J.A., S. Shalini, J.M. Risk, C.L. Day, and S. Kumar. 2010. A direct interaction with NEDD1 regulates gamma-tubulin recruitment to the centrosome. PLoS One. 5:e9618.

Muroyama, A., L. Seldin, and T. Lechler. 2016. Divergent regulation of functionally distinct γ-tubulin complexes during differentiation. J. Cell Biol. 213:679–692.

Pinyol, R., J. Scrofani, and I. Vernos. 2013. The role of NEDD1 phosphorylation by Aurora A in chromosomal microtubule nucleation and spindle function. Curr. Biol. 23:143–149.

Punjani, A., J.L. Rubinstein, D.J. Fleet, and M.A. Brubaker. 2017. cryoSPARC: algorithms for rapid unsupervised cryo-EM structure determination. Nat. Methods. 14:290–296.

Sánchez-Huertas, C., F. Freixo, R. Viais, C. Lacasa, E. Soriano, and J. Lüders. 2016. Non-centrosomal nucleation mediated by augmin organizes microtubules in post-mitotic neurons and controls axonal microtubule polarity. Nat. Commun. 7:12187.

Sdelci, S., M. Schütz, R. Pinyol, M.T. Bertran, L. Regué, C. Caelles, I. Vernos, and J. Roig. 2012. Nek9 phosphorylation of NEDD1/GCP-WD contributes to Plk1 control of γ-tubulin recruitment to the mitotic centrosome. Curr. Biol. 22:1516–1523.

Serna, M., F. Zimmermann, C. Vineethakumari, N. Gonzalez-Rodriguez, O. Llorca, and J. Lüders. 2024. CDK5RAP2 activates microtubule nucleator γTuRC by facilitating template formation and actin release. Dev. Cell. 0. doi:10.1016/j.devcel.2024.09.001.

Ti, S.-C., M. Wieczorek, and T.M. Kapoor. 2020. Purification of Affinity Tag-free Recombinant Tubulin from Insect Cells. STAR Protoc. 1. doi:10.1016/j.xpro.2019.100011.

Tovey, C.A., and P.T. Conduit. 2018. Microtubule nucleation by γ-tubulin complexes and beyond. Essays Biochem. 62:765–780.

Uehara, R., R.-S. Nozawa, A. Tomioka, S. Petry, R.D. Vale, C. Obuse, and G. Goshima. 2009. The augmin complex plays a critical role in spindle microtubule generation for mitotic progression and cytokinesis in human cells. Proc. Natl. Acad. Sci. U. S. A. 106:6998–7003.

UniProt Consortium. 2023. UniProt: The universal protein knowledgebase in 2023. Nucleic Acids Res. 51:D523–D531.

Viais, R., M. Fariña-Mosquera, M. Villamor-Payà, S. Watanabe, L. Palenzuela, C. Lacasa, and J. Lüders. 2021. Augmin deficiency in neural stem cells causes p53-dependent apoptosis and aborts brain development. Elife. 10. doi:10.7554/eLife.67989.

Waterhouse, A.M., J.B. Procter, D.M.A. Martin, M. Clamp, and G.J. Barton. 2009. Jalview Version 2--a multiple sequence alignment editor and analysis workbench. Bioinformatics. 25:1189–1191.

Wieczorek, M., T.-L. Huang, L. Urnavicius, K.-C. Hsia, and T.M. Kapoor. 2020a. MZT Proteins Form Multi-Faceted Structural Modules in the γ-Tubulin Ring Complex. Cell Rep. 31:107791.

Wieczorek, M., S.-C. Ti, L. Urnavicius, K.R. Molloy, A. Aher, B.T. Chait, and T.M. Kapoor. 2021. Biochemical reconstitutions reveal principles of human γ-TuRC assembly and function. J. Cell Biol. 220. doi:10.1083/jcb.202009146.

Wieczorek, M., L. Urnavicius, S.-C. Ti, K.R. Molloy, B.T. Chait, and T.M. Kapoor. 2020b. Asymmetric Molecular Architecture of the Human γ-Tubulin Ring Complex. Cell. 180:165–175.e16.

Würtz, M., E. Zupa, E.S. Atorino, A. Neuner, A. Böhler, A.S. Rahadian, B.J.A. Vermeulen, G. Tonon, S. Eustermann, E. Schiebel, and S. Pfeffer. 2022. Modular assembly of the principal microtubule nucleator γ-TuRC. Nat. Commun. 13:473.

Xu, Y., H. Muñoz-Hernández, R. Krutyhołowa, F. Marxer, F. Cetin, and M. Wieczorek. 2024. Partial closure of the γ-tubulin ring complex by CDK5RAP2 activates microtubule nucleation. Dev. Cell. 0. doi:10.1016/j.devcel.2024.09.002.

Zeng, T., Y. Lee, and B. Liu. 2009. The WD40 repeat protein NEDD1 functions in microtubule organization during cell division in Arabidopsis thaliana[W]. The Plant Cell Online. 21:1129–1140.

Zhang, J., I.R. Humphreys, J. Pei, J. Kim, C. Choi, R. Yuan, J. Durham, S. Liu, H.-J. Choi, M. Baek, D. Baker, and Q. Cong. 2024. Computing the Human Interactome. bioRxiv. doi:10.1101/2024.10.01.615885.

Zhang, X., Q. Chen, J. Feng, J. Hou, F. Yang, J. Liu, Q. Jiang, and C. Zhang. 2009. Sequential phosphorylation of Nedd1 by Cdk1 and Plk1 is required for targeting of the gammaTuRC to the centrosome. J. Cell Sci. 122:2240–2251.

Zhang, Y., X. Hong, S. Hua, and K. Jiang. 2022. Reconstitution and mechanistic dissection of the human microtubule branching machinery. J. Cell Biol. 221. doi:10.1083/jcb.202109053.

Zhong, E.D., T. Bepler, B. Berger, and J.H. Davis. 2021. CryoDRGN: reconstruction of heterogeneous cryo-EM structures using neural networks. Nat. Methods. 18:176–185.

Zhuo, T., Z. Wu, S. Chen, C. Yang, H. Huang, J. Gan, J. Lyu, J. Xiao, Z. Li, S. Qin, and Y. Wu. 2024. NEDD1 overexpression increases cell proliferation, tumor immune escape, and drug resistance in LUAD. J. Cancer. 15:2460–2474.

Zimmermann, F., M. Serna, A. Ezquerra, R. Fernandez-Leiro, O. Llorca, and J. Luders. 2020. Assembly of the asymmetric human γ-tubulin ring complex by RUVBL1-RUVBL2 AAA ATPase. Science Advances. doi:10.1126/sciadv.abe0894.

